# Whole-genome sequencing confirms multiple species of Galapagos giant tortoises

**DOI:** 10.1101/2023.04.05.535692

**Authors:** Stephen J. Gaughran, Rachel Gray, Menna Jones, Nicole Fusco, Alexander Ochoa, Joshua M. Miller, Nikos Poulakakis, Kevin de Queiroz, Adalgisa Caccone, Evelyn L. Jensen

## Abstract

Galapagos giant tortoises are endemic to the Galapagos Archipelago, where they are found in isolated populations. While these populations are widely considered distinguishable in morphology, behavior, and genetics, the recent divergence of these taxa has made their status as species controversial. Here, we apply multispecies coalescent methods for species delimitation to whole genome resequencing data from 38 tortoises across all 13 extant taxa to assess support for delimiting these taxa as species. In contrast to previous studies based solely on divergence time, we find strong evidence to reject the hypothesis that all Galapagos giant tortoises belong to a single species. Instead, a conservative interpretation of model-based and divergence-based results indicates that these taxa form a species complex consisting of a minimum of 9 species, with some analyses supporting as many as 13 species. There is mixed support for the species status of taxa living on the same island, with some methods delimiting them as separate species and others suggesting multiple populations of a single species per island. These results make clear that Galapagos giant tortoise taxa represent different stages in the process of speciation, with some taxa further along in that evolutionary process than others. A better understanding of the more complex parts of that process is urgently needed, given the threatened status of Galapagos giant tortoises.

**Lay Summary:** Species delimitation is a challenging problem in evolutionary biology, but one that is central to the field. Distinguishing species can affect conservation management practices, from conservation status assessments to strategies for breeding programs. More fundamentally, understanding species boundaries affects our ability to assess biodiversity and to study evolutionary processes. The Galapagos Archipelago presents several radiations of closely related taxa that inspired Charles Darwin to develop his theory of evolution by natural selection and later led to foundational case studies in speciation. The Galapagos giant tortoises were one such inspiration. Nearly two centuries later, there is still an ongoing debate about the taxonomic status of these tortoises, with opinions on their status ranging from barely differentiated populations to separate species. Here, we present the first genomic species delimitation of Galapagos giant tortoises and provide convincing evidence that this group is a complex consisting of between 9 and 13 species. These results provide valuable guidance to conservation stakeholders in the Galapagos, while also adding an important case study to the delimitation of island species.

## Introduction

Speciation is a complex biological process driven at least in part by ecological context such as physical barriers to gene flow, adaptation to local environments, and population-specific demographic dynamics. In recently diverged lineages, the relative roles of the evolutionary forces responsible for divergence can be challenging to describe accurately because of shared ancestral polymorphisms in the descendant lineages and introgression due to ongoing gene flow (Shaffer & Thomson 2007). Adding to these difficulties are the small founding populations that characterize the origin of some species, especially in island settings, which can speed up the divergence of lineages through the rapid loss or fixation of alleles (Kimura & Ohta 1969). The iconic radiation of Galapagos giant tortoises (a clade within the genus *Chelonoidis*) is a compelling example of the complexities of species delimitation in a case of recent diversification: their divergence has been molded by a combination of vicariance and colonization events (Caccone *et al*. 1999, 2002; Poulakakis *et al*. 2012, 2020) brought about by both natural and anthropogenic environmental changes and making accurate species delimitation challenging.

The Galapagos Islands, the conceptual home of the theory of evolution by natural selection, provide a crucible for the study of speciation and taxonomic complexity. Within the Galapagos Archipelago there are many examples that show clear evidence of recently diverged but genetically distinct species, including finches (Grant & Grant 2003), lava lizards (Benavides *et al*. 2009), iguanas (MacLeod *et al*. 2015), mockingbirds (Arbogast *et al*. 2006), moths (Schmitz *et al*. 2007), and tortoises (Caccone *et al*. 1999). This archipelago therefore provides a model system to understand the complex realities of speciation, allowing us to explore multiple lines of molecular, morphological, and ecological evidence when proposing species delimitations.

There are 16 Galapagos giant tortoise taxa, three of which (*niger*, *abingdonii*, and an unnamed taxon from Santa Fé Island) are recently extinct (Figure 1). Morphological, behavioral, and genetic differences have been documented among the taxa (Gaughran *et al*. 2018, Jensen *et al*. 2021, Chiari 2021, Hunter *et al*. 2013). The most striking morphological difference is carapace shape, which ranges from a domed shape to a saddleback shape with an elevated anterior carapace opening. These carapace shapes have a genetic underpinning, as evidenced by a clear phylogenetic signal (Jensen *et al*. 2022) and a strong inheritance in juveniles of different taxa raised in a common environment (Pritchard 1996). The taxa also differ in coloration on the head and neck, as well as in limb length (reviewed in Chiari 2021) and aggressive behaviors (Schafer & Krekorian 1983). Mitochondrial, microsatellite, and genomic data all suggest that geographically distinct populations represent genetically differentiated lineages (e.g., Caccone *et al*. 2002, Beheregaray *et al*. 2003, Gaughran *et al*. 2018, Miller *et al*. 2018).

**Figure 1.**
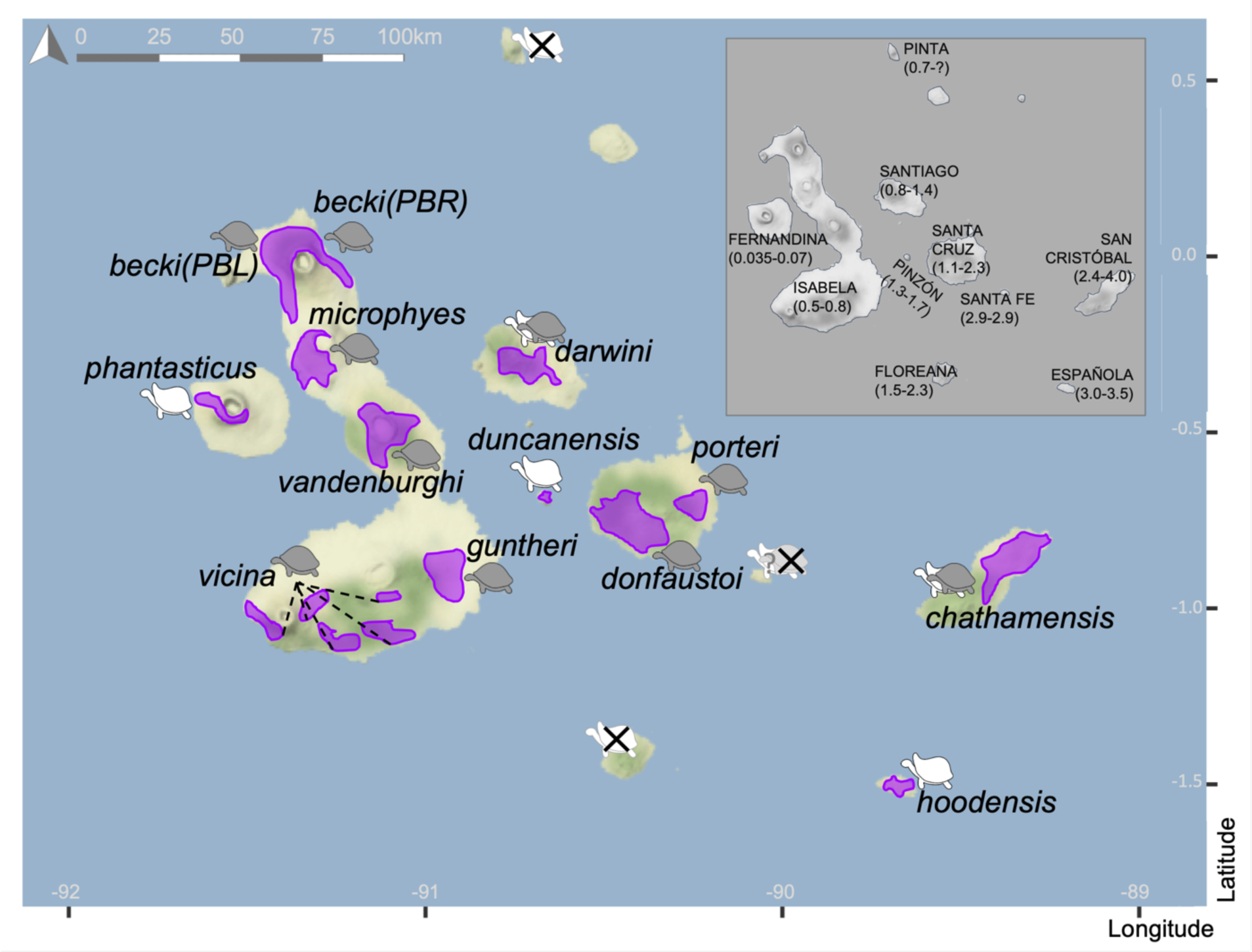
Galapagos giant tortoises are found on seven islands and form at least 13 genetically distinct taxa. Purple shapes indicate current ranges of populations. Carapace morphology (domed = gray, saddleback = white) is shown for each population, with “intermediate” shell shape indicated with overlapping icons. Icons with an “X” indicate extinct tortoise populations. Isabela Island and Santa Cruz Island are both home to multiple genetically distinct, allopatric populations. Inset map shows the names of islands with recently extant Galapagos giant tortoise taxa, with geological ages of each island in million years below.

Galapagos giant tortoises are descended from individuals that likely floated from continental South America along the Humboldt Current to the newly formed Galapagos Islands around 3 million years ago (Ma; Caccone *et al*. 2002). As more volcanic islands emerged over time, tortoises migrated from existing islands in the east to newer ones in the west (Poulakakis *et al*. 2020). Divergence dates from mitochondrial DNA suggest that the progressive colonization from the first colonized island likely occurred after 2 Ma and continued up to the emergence of the youngest island to the west, Fernandina, around 60,000 years ago (Geist *et al*. 2014). The layout of the archipelago has changed dramatically over this time period due to volcanic emergence and subsidence, and sea level changes. During the Pleistocene the central archipelago had a larger landmass that has since fragmented into some of the islands seen today (Geist *et al*. 2014). Thus, the diversification of Galapagos giant tortoises has been triggered by a combination of dispersal to isolated islands and vicariance as landmasses fragmented (Parent *et al*. 2008). In addition, within the last 200 years, humans have moved tortoises between islands (Caccone *et al*. 2002, Poulakakis *et al*. 2008) and land use change has brought previously isolated lineages into contact (Russello *et al*. 2005), resulting in low levels of recent gene flow between previously isolated populations.

Because of this recent and dynamic evolutionary background, the taxonomy of the Galapagos giant tortoises has been the subject of almost endless debate since the time of Darwin (e.g., Darwin 1839, Günther 1877, Van Denburgh 1907, Pritchard 1996, Zug 1998, Caccone *et al*. 1999, Kehlmaier *et al*. 2021). Most taxonomic proposals have recognized these lineages as either a single species with 2-14 subspecies, or as many as 14 separate species (reviewed in Pritchard 1996). Following work by Caccone *et al*. (1999), they have been listed as multiple species based on genetic data accumulated over the last several decades (Rhodin *et al*. 2017). More recently, however, two publications have advocated collapsing all taxa into a single species with multiple subspecies (Loire *et al*. 2013, Kehlmaier *et al*. 2021), which was adopted by the IUCN Tortoise and Freshwater Turtle Specialist Group in 2021 (Rhodin *et al*. 2021).

To address this question, we turned to whole genome resequencing data and methods that quantitatively assess the degree to which taxa are independently evolving under a multispecies coalescent (MSC) framework (Zhang *et al*. 2011, Jackson *et al*. 2017, Morales *et al*. 2018, Mays *et al*. 2019, Leaché *et al*. 2019, Marshall *et al*. 2021). Here, we apply several of these methods of species delimitation to the Galapagos giant tortoises, and thereby provide an assessment of taxonomic considerations in this endangered clade.

## Methods

### Terminology, study design and data

The aim of this paper is to assess genomic evidence for the distinctiveness and taxonomic status of Galapagos giant tortoise taxa using phylogenetic and coalescent frameworks. To remain agnostic about the taxonomic status of the various tortoise populations prior to presenting our results, we avoid designating the taxa as *species* or *subspecies* and refer to them using only the epithets (e.g., *phantasticus* rather than *Chelonoidis phantasticus* or *Chelonoidis niger phantasticus*). We mapped existing Illumina short-read data (NCBI Bioproject PRJNA761229; Jensen *et al*. 2021, Jensen *et al*. 2022) to the Pinta Island Galapagos giant tortoise reference genome (NCBI assembly ASM359739v1, Quesada *et al*. 2019). Samples in this short-read data set included two individuals of *phantasticus*, and three individuals each of *guntheri*, *microphyes*, *vandenburghi*, *vicina*, *becki* (PBL population), *becki* (PBR population), *darwini*, *chathamensis*, *donfaustoi*, *porteri*, *duncanensis*, and *hoodensis*, for a total of 38 Galapagos giant tortoise individuals plus one outgroup (Chaco tortoise, *C. chilensis*) individual. We called variant sites for each individual using the BCFtools variant calling pipeline (Li *et al*. 2009) and created a consensus fasta file for each individual from the VCF. We then generated a random set of 1kb loci separated by 100kb. We filtered, masked, and generated phased haplotypes for each locus. Details on these terminology choices, study design, and data are in the Supplemental Methods.

### SNAPP tree generation

To assess the phylogenetic relationships among the extant populations of Galapagos giant tortoises, we used SNAPP v1.5.2 (Bryant *et al*. 2012) and MODEL_SELECTION v1.5.3 (Baele *et al*. 2012), as implemented in BEAST2 v2.6.7 (Bouckaert *et al*. 2014), to test 21 different phylogenetic models (Supplemental Table S1) on a dataset comprising 1,000 SNPs from 38 Galapagos giant tortoise individuals, plus the *C. chilensis* individual as an outgroup. We then applied the priors from the best model to an extended dataset comprising 5,000 SNPs for phylogenetic reconstruction and downstream species delimitation comparisons. A more detailed description of these methods is presented in Supplemental Methods.

### Species delimitation models

We implemented the Bayesian species delimitation model in BPP v4.0 (Flouri *et al*. 2018) to test hypotheses about the number of species in our data set. We ran the species delimitation analyses with 50, 200, 500, or 1000 phased loci, with each locus a randomly selected alignment of 1000 bp (see Supplemental Methods). We ran BPP species delimitation with the SNAPP guide tree (i.e., “A10” analysis) and without a guide tree (i.e., “A11” analysis), repeating each run three times with different random seeds to assess convergence among runs. Because the Galapagos Islands are geologically young (<3.5 Ma) and the deepest divergence time among Galapagos giant tortoise lineages is estimated to be between 2 and 3 Ma (Caccone *et al*. 2002), we chose a small but diffuse prior for divergence time (*τ* ∼ IG(3, 0.001)), which corresponds to a mean of 83,333 generations (approximately 2.08 million years). In addition, given the low estimates of effective population size for Galapagos giant tortoises (Garrick *et al*. 2015, Jensen *et al*. 2021) and because the carrying capacity must be low for large-bodied terrestrial vertebrates on these semi-desert islands, we used a prior scaled effective population size of *θ* ∼ IG(3, 0.001), which is equivalent to *N_e_*=20,833. However, because there is uncertainty about historical population size, we also re-ran the analyses with scaled effective population size priors that were larger (*θ* ∼ IG(3, 0.005), *N_e_*=104,167) or smaller (*θ* ∼ IG(3, 0.0001), *N_e_*=2083), keeping the priors for τ the same. In each of run, we used a burn-in of 100,000, a sampling frequency of 2, and collected 300,000 samples.

To assess if the chosen loci and included taxa affected our analyses, we ran the above analyses with a different set of 50, 200, 500, and 830 loci and including the Chaco tortoise. The inclusion of this outgroup allowed the model to explore a species delimitation of two (i.e., Galapagos giant tortoises and Chaco tortoise). Because the divergence time estimate of the Chaco tortoise and Galapagos giant tortoises is around 12 Ma (Caccone *et al*. 1999, Kehlmaier *et al*. 2017), we used a divergence time prior of *τ* ∼ IG(3, 0.01). Finally, we re-ran the analyses with 50 unphased loci to ensure that our phasing method was not affecting the results.

### Genealogical divergence index (*gdi*) from BPP

The use of species delimitation models in BPP is thought to be a robust method for sympatric species delimitation, but there are concerns that using this MSC-based model selection for species delimitation may delimit populations rather than species, especially if the taxa are allopatric (Jackson *et al*. 2017, Leaché *et al*. 2019). Therefore, we also calculated the *gdi*, a distance-based method with heuristic cutoffs for species delimitation (Jackson *et al*. 2017). The *gdi* is calculated from the divergence time in coalescent units (2*τ/θ*) and reflects the probability that two sequences from a purported species coalesce before the divergence time with a sister species (*τ*) (Leaché *et al*. 2019). As such, it is a coalescent-based measure of genetic divergence, as opposed to a sequence-based measure. Using the effective population size and divergence time, the *gdi* can be calculated in a pairwise manner for sister species *a* and *b*, and can range from 0 (panmixia) to 1 (strong divergence). Based on empirical data and simulations, Jackson *et al*. (2017) proposed a general heuristic for delimiting species based on the *gdi*. Namely, *gdi* < 0.2 indicates a single species, and *gdi* > 0.7 indicates different species. Values for *gdi* between these cutoffs indicate ambiguous delimitation.

To calculate these indices, we used BPP for parameter estimation of *τ* and *θ*, under the guide tree and fixed MSC model (“A00”) function in BPP. For these analyses, we used the 830 phased loci available in the data set with the Chaco outgroup. Using the phylogeny from our SNAPP analysis as a fixed guide tree, we estimated *τ* for each node and *θ* for each branch and tip, by running the MCMC with a burn-in of 100,000, a sample frequency of 2, and a total sample of 200,000. We used the same loci and priors as in the species delimitation modeling. We then used the MCMC output to create posterior distributions of *τ* (for each node) and *θ* (for each tip), which we then used to calculate *gdi*. Using posterior distributions allowed us to calculate 95% credibility intervals. Following the suggestion of Leaché *et al*. 2019, we also successively collapsed each node of the phylogeny, re-ran the A00 parameter estimation model, and calculated *gdi* for each collapsed taxon.

### PHRAPL

We then assessed the presence of gene flow between all pairs of lineages within each of the two main clades of the tree (domed and saddleback) and tested for a collapse event using PHRAPL (Jackson *et al*. 2017) implemented in R version 4.2 (R Core Team, 2022). PHRAPL works by estimating the probability that a set of gene trees are observed under a given model by calculating the frequency at which they are observed in a distribution of expected tree topologies, weighting the probability by Akaike information criterion (AIC). In this way, PHRAPL can determine the most likely demographic history. Three demographic models for each pairwise comparison were constructed to test varying divergence and gene flow scenarios between populations of Galapagos giant tortoises (Supplemental Figure S1). Model 1 was a two species isolation only model, which tested for divergence between two populations with no ongoing migration. Model 2 was also a two-species model and tested for constant symmetrical migration between the two populations. The final model (Model 3) was the same as Model 2 but we removed the divergence between the two populations using the *setCollapseZero* function in PHRAPL. This single species model therefore allowed for constant symmetrical gene flow between the populations with no divergence events. Input gene trees were generated for each of the 830 phased loci in RaxML (v 8.2.12, Stamatakis 2014) with 20 replicate searches, rapid hill-climbing and the GTRGAMMA model. The Chaco tortoise was included as an outgroup to root the tree. PHRAPL is a demographic model and requires individuals to be assigned to a population prior to model construction. Here, individuals were assigned to a population based on geographic location, as detailed in Figure 1. In this way, the tortoises on each island were represented by a single population except that six geographically separate populations were recognized on Isabela Island and two on Santa Cruz Island. each island lineage was assigned to a distinct population, with the exception of Isabela Island having six populations, based on the six geographically separated lineages, and Santa Cruz island having two populations. Gene trees were subsampled at random with replacement 100 times, sampling 2 individuals per lineage in each replicate. The outgroup was not included in the subsampling and so was not included in the PHRAPL runs. Simulation of 100,000 gene trees was conducted using a grid of parameter values for divergence time (*t*) and migration (*m*; Supplemental Table S2). The initial divergence within the Galapagos tortoise radiation is estimated to be within the last 2 Ma, but the species pairs we compare here are more recent (Poulakakis *et al*. 2020). We set the parameters of our grid search to capture this by limiting the maximum divergence (*t*) to 1 Ma. Migration rates were equal to 4*Nm*, where *Nm* is the number of migrants per generation, with the lowest migration being equivalent to one migrant every 800 years and the highest equivalent to one migrant every 25 years. This range of values was designed to capture the putatively complex historical gene flow among populations. Akaike weights (wAIC) were used to compare models and calculate model probabilities ranging from 0 (low support) to 1 (high support). To present the best supported hypothesis overall for each pairwise comparison, the summed wAIC of the two-species models (Models 1 and 2) was compared with the wAIC of the single species model (Model 3). We interpreted a summed wAIC of > 0.9 across the two-species models as strong support for a two species hypothesis, a wAIC > 0.9 of the one species model as strong support for a single species hypothesis, and all other scenarios as ambiguous. In addition to identifying the top model chosen by PHRAPL, we also calculated the *gdi* value between taxon pairs following the approach taken in Jackson *et al*. 2017 and using the *CalculateGdi* function in PHRAPL. The *gdi* value from PHRAPL differs from that from BPP in that it is calculated using the model averaged divergence rate (*t*), migration rate into population 1 (*M1*), and migration into population 2 (*M2*). Because our models did not estimate the direction of migration, *M1=M2* in our analyses. We interpreted the *gdi* values using the same thresholds described above.

## Results

### SNAPP

Of the 21 SNAPP models tested, Model 6 (α = 5, β = 150, λ = 39) presented the greatest marginal likelihood estimate (–14,448.23), with a Bayes factor equivalent to –299.07 relative to the default, reference model (*i.e.*, Model 7; Table S1). Using the framework of Kass & Raftery (1995), the Bayes factor in support for Model 6 is “decisive” compared to the reference model. The final tree derived from the parameters used for Model 6 (combined posterior effective sample size = 1,085; cladogram shown in Figure 2, phylogram with scaled branch lengths in Supplemental Figure S2) has two primary sub-clades, which reflect the two carapace morphologies: the larger clade contains the nine taxa with mostly domed carapaces, whilst the smaller clade contains the four taxa with mostly saddleback carapaces. Most nodes (82%) are highly supported, with posterior probabilities ≥ 0.95 (Figure 2).

**Figure 2.**
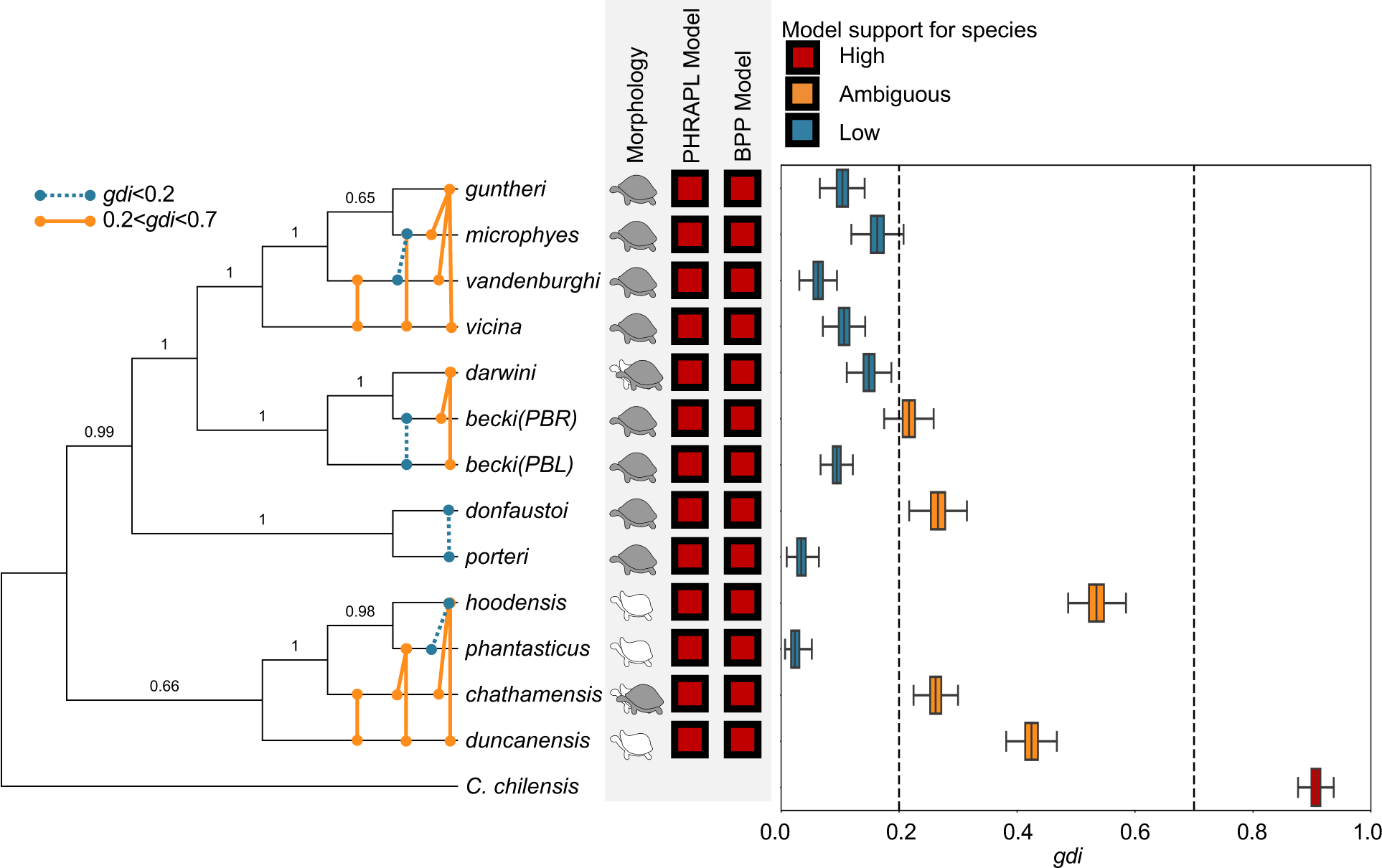
Multiple lines of evidence confirm the Galapagos giant tortoise species complex consists of multiple genetically divergent, independently evolving species. On the left, the cladogram generated from genome-wide SNPs in SNAPP, with branch posterior support values. Within-clade PHRAPL pairwise *gdi* values are indicated on the tree by colored lines connecting lineages. In the center, support for species delimitation models in PHRAPL and BPP. On the right, a boxplot of the *gdi* values estimated from the posterior distributions of *τ* and *θ* generated by BPP for each Galapagos giant tortoise taxon (red: *gdi* > 0.7, meeting the heuristic threshold for strongly delimited species; blue: *gdi* < 0.2, below the heuristic threshold for a single species; orange: 0.2 < *gdi* < 0.7, which is considered ambiguous species delimitation). Model-based species delimitation in BPP and PHRAPL model selection supported 13 species (not shown).

### Species delimitation models in BPP

When we ran an “unguided” species delimitation (A11 analysis in BPP) with our most realistic prior of *θ* ∼ IG(3, 0.001), the analysis supported 13 species with P>0.98 when using 50, 200, 500, or 1000 phased loci across all runs (Supplemental Table S3). When a smaller *θ* prior was used, 13 species were almost always supported with P>0.99 (Supplemental Table S3). When using a slightly larger *θ* prior, the analysis generally supported 12 or 13 species when 200, 500, and 1000 loci were used, but as few as 9 species when only 50 loci were used (Supplemental Table S3). In cases where there was support for fewer than 13 species, the analysis typically supported the collapse of two or more taxa on Isabela Island into one species. In four out of 45 runs, the A11 analysis supported a single species of Galapagos giant tortoise (Supplemental Tables S3 and S5), but these runs likely represent poor mixing (see Discussion). When the Chaco tortoise outgroup was included, most runs supported more than 10 species (Supplemental Table S4). Importantly, a two-species model (with Galapagos giant tortoises as one species and the Chaco tortoise as another species) was never supported. However, runs with the Chaco tortoise did not always converge on the same distributions of posterior support when started with different random seeds (Supplemental Table S4). Results were nearly identical when we used the unphased loci (Supplemental Tables S5-S6) and running the “A10” analysis with the guide tree (Supplemental Tables S7-S8). Using our most realistic priors, therefore, the BPP analyses supported delimitation of 13 species (Figure 2).

### *gdi* calculation from BPP parameter estimates

Overall, the median *gdi* estimated in BPP ranged from 0.023 (*phantasticus*) to 0.534 (*hoodensis*; Figure 2, Table S11). None of the taxa exceeded a *gdi* of 0.7, which has been proposed as a heuristic for strongly delimited species. Five taxa (*becki (PBR)*, *donfaustoi*, *hoodensis*, *duncanensis*, and *chathamensis*) had *gdi* in the ambiguous delimitation range of 0.2 < *gdi* < 0.7. The remaining eight taxa had *gdi* below 0.2. Estimates of *gdi* were effectively identical across priors (Supplemental Tables S12–S13). In many cases, the *gdi* differed dramatically between sister taxa, emphasizing the asymmetry commonly observed in this statistic.

When taxa do not meet the heuristic *gdi* threshold for species delimitation, Leaché *et al*. (2019) recommended progressively collapsing taxon pairs, rerunning the MCMC and calculating the *gdi* of the new groups. If the collapsed taxon represents a better-supported species, the expectation is that the *gdi* would increase. This occurred when the taxa on central and southern Isabela Island were collapsed (*gdi*=0.226, Figure 3A and Supplemental Table S16) and when those on Santa Cruz Island were collapsed (*gdi*=0.291, Figure 3C and Supplemental Table S14). In all other cases, however, the collapsed taxa had *gdi* lower than the separated taxa (Figure 3B and 3D, Supplemental Tables S14-S18). Overall, the *gdi* estimates provide ambiguous support for delimiting most taxa on different islands as separate species but favor treating some populations on the same island as conspecific (Figure 2).

**Figure 3.**
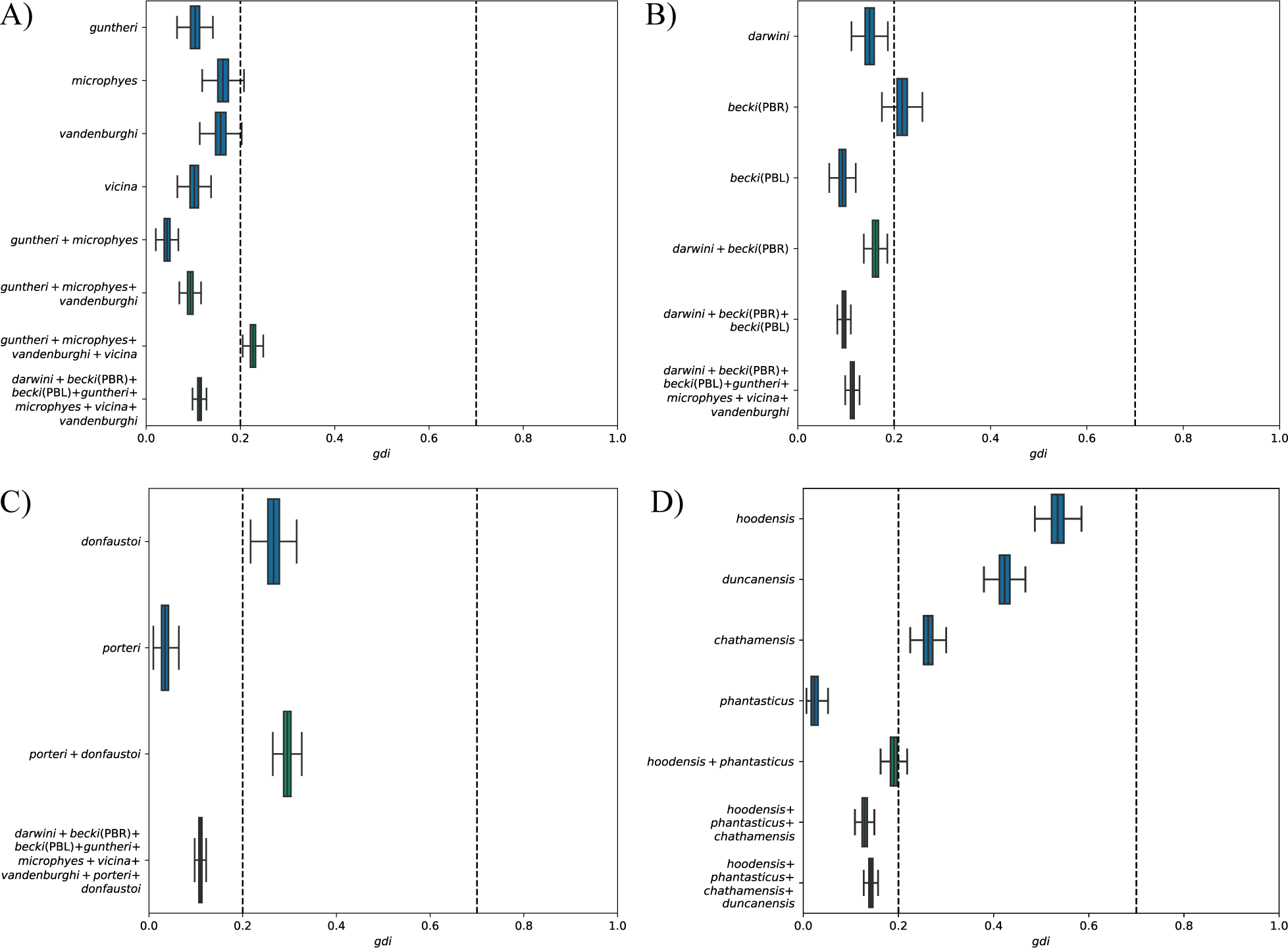
Estimate *gdi* for progressively collapsed taxa shows support for some delimited species but not others. *gdi* calculated for each taxon, including when sister taxa are collapsed following the method of Leaché *et al*. (2019). Blue boxes are single named taxa; green boxes are two or more taxa collapsed into a single “species.” A) Isabela Island clade with progressive taxonomic collapse, showing a higher *gdi* above 0.2 when the four taxa from central and southern Isabela Island are collapsed. B) Santiago Island and northern Isabela Island clade with progressive taxonomic collapse, showing lower *gdi* when taxa are collapsed. C) Santa Cruz Island clade with progressive taxonomic collapse, showing a higher *gdi* above 0.2 when the two taxa are collapsed. D) Saddleback clade with progressive taxonomic collapse, showing lower *gdi* when taxa are collapsed.

### PHRAPL

For the PHRAPL analysis we focus on results of pairwise comparisons for populations within the two primary clades (Figure 2). For the saddleback clade, there are four lineages spread across four separate islands and so all pairwise comparisons were included here. Within the domed clade, Isabela and Santa Cruz islands host five and two lineages respectively, while Santiago has a single living lineage. For this clade we have presented the results in Figure 2 as within Santa Cruz Island for its two lineages, within central and southern Isabela Island for its four lineages that form a clade and within the clade of Santiago and its sister taxa, *becki (PBR)* and *becki (PBL)* on northern Isabela Island. Results for model wAIC and *gdi* values for all possible pairwise comparisons within the saddleback and domed morphology clades are presented in Supplemental Table S20. Within the saddleback clade, all pairwise comparisons supported a two-species model when wAIC values were summed, supporting four species across this clade. The *gdi* values in this clade were generally in the ambiguous range of 0.2<*gdi*<0.7, except for the *gdi* of *hoodensis* and *phantasticus* (*gdi*=0.179; Figure 2 and Table S20). For the Santa Cruz Island tortoises (*porteri* and *donfaustoi*), PHRAPL supported a two-species model with a wAIC > 0.9, but the *gdi* was 0.181 (Figure 2). Similarly, within the central and southern Isabela Island clade there was full support for two-species models in pairwise comparisons, yet mixed support from the *gdi* values across pairwise comparisons of the four lineages. The pairwise comparisons between taxa on Santa Cruz and Isabela Islands also all favored a two-species model, and all *gdi* estimates fell between 0.2 and 0.7 (Figure 2). Similarly, all comparisons between the Santiago lineage (*darwini*) and the taxa on Isabela and Santa Cruz Islands strongly supported a two-species model, with five out of eight *gdi* comparisons above 0.2. In total, the species delimitation models in PHRAPL support the delimitation of 13 species, while the *gdi* estimates from PHRAPL suggest between 7 and 9 species, depending on the interpretation of heuristic thresholds.

## Discussion

Despite considerable progress in achieving a more unified concept of species (Mayden 1997; de Queiroz 1998, 2007; Hey 2006), the empirical application of such a concept presents challenges, particularly in cases of recent and/or incomplete divergence (e.g., de Queiroz 2005a, Carstens *et al*. 2013, Jackson *et al*. 2017). The Galapagos giant tortoises represent such a case. Here, we build on decades of research on the molecular evolution of these tortoises and provide genomic evidence that the Galapagos giant tortoise complex consists of multiple distinct species.

Our phylogenetic analysis using SNAPP supported the topology previously inferred from genome-wide sequence data (Jensen *et al*. 2022), except for the relationships among the most recently diverged taxa (*guntheri*, *vandenburghi* and *microphyes*) and the placement of the Santa Cruz Island taxa. Given this strong support, we used this nuclear phylogeny as a guide tree in subsequent analyses. Notably, the topology of these trees inferred from nuclear loci deviates in some ways from mitochondrial trees for these taxa (Poulakakis *et al*. 2012, 2020), an incongruence that highlights the complex and rapid evolutionary history of Galapagos giant tortoises.

Our first line of evidence for the distinctiveness of Galapagos giant tortoise species is the model-based species delimitation analysis in BPP. This analysis supported 13 species of Galapagos giant tortoises across different numbers of loci when our most realistic priors were applied. Across different priors we observe stronger support for 13 species as more loci are used, which may indicate that the greater information content from more loci provides better biological resolution. Theory and simulations suggest that BPP will increasingly favor a two-species model when hundreds or thousands of loci are used, even under high rates of migration (Leaché *et al*. 2019). Under our most realistic prior set, however, even 50 loci strongly support the delimitation of 13 species both with and without a guide tree.

In cases where fewer than 13 species were supported, the model favored collapsing two or more of the taxa in a clade of four taxa on Isabela Island (i.e., *guntheri*, *microphyes*, *vandenburghi*, and *vicina*). These taxa are geographically adjacent to each other on the central and southern volcanoes of Isabela Island, which suggests that recent divergence or gene flow is a factor. The delimitation models sometimes collapsed taxa that were not sister taxa on our phylogeny, emphasizing the finding from this and other studies that the phylogeny of taxa on Isabela Island is complex (Poulakakis *et al*. 2012, Jensen *et al*. 2022) and likely needs genomic data from a larger number of individuals to be resolved.

In four runs of our A11 analysis including only Galapagos giant tortoise individuals the analysis supported a single species of Galapagos giant tortoise. These runs are likely the result of poor mixing, which is a known issue in the reversible-jump Markov Chain Monte Carlo (rjMCMC) algorithm of BPP (Yang and Rannala 2010, Flouri *et al*. 2018). Mixing can be especially problematic when sampling the fully resolved or fully collapsed tree, where the algorithm can get “stuck” (Giarla *et al*. 2014). That we never find support for a single Galapagos giant tortoise species when the Chaco tortoise individual is added to the analysis nor when we provide a guide tree further supports the conclusion that the single-species runs of the A11 analysis are the result of poor mixing.

We also modeled species delimitation in PHRAPL, comparing support for models of each taxonomic pair as one species, two species in isolation, and two species with gene flow. We found two-species models favored in all cases (Figure 2, Supplemental Table S20). The migration rates chosen accounted for migration occurring up to 800 generations ago and there was mixed support across populations for the two-species model that allowed migration vs. no migration. Jackson *et al*. (2017) described the process of model selection in PHRAPL to be less accurate for isolation only models (here, Model 1) with recent divergence times (*t*<2). Given that we tailored the divergence values to be <1 to suit the evolutionary time scale of the Galapagos giant tortoise lineages, there may be some bias in results where an isolation-with-migration model is selected over the isolation-only model. This is a limitation of PHRAPL in that it has difficulty distinguishing incomplete lineage sorting from gene flow, and so will favor the isolation-with-migration model over the model of recent isolation. Furthermore, PHRAPL is more accurate with a larger dataset (Jackson *et al*. 2017). The sample size for each taxon in our data set is small (n=2– 3 individuals per taxon). Because PHRAPL uses subsampling of individuals to generate expected tree topologies, having a larger sample size that better represents the whole population could help disentangle the history of the unresolved populations. Current work is underway to increase the number of reference sequences for extant populations.

The species delimitation model in BPP has been criticized as oversplitting species and recovering population structure rather than species (Jackson *et al*. 2017), as has species delimitation modeling in PHRAPL (Leaché *et al*. 2019). Some have argued that the *gdi* better reflects species differences by explicitly incorporating information about divergence time and either population size (when calculated from BPP parameter estimates) or migration rates (when calculated in PHRAPL) (Jackson *et al*. 2017, Leaché *et al*. 2019). Using the *gdi* thresholds of *gdi*<0.2 for single species and *gdi*>0.7 for two species, we found ambiguous support (0.2<*gdi*<0.7) in the BPP *gdi* estimates for *becki (PBR)*, *donfaustoi*, *hoodensis*, *chathamensis*, and *duncanensis*. All other taxa had a *gdi*<0.2, creating multiple scenarios in which one sister taxon had *gdi*>0.2 and the other taxon had *gdi*<0.2. In PHRAPL, the *gdi* estimates for most taxa were also in the ambiguous range of 0.2<*gdi*<0.7, with some taxon pairs falling below 0.2 (i.e., *phantasticus* and *hoodensis*, *porteri* and *donfaustoi*, and some of the domed taxa on Isabela and Santiago Islands).

Although these thresholds were proposed in the literature from a small sample of species, work in other natural populations has shown that well-accepted species can fall into the ambiguous range of 0.2–0.7, including horned lizards (Leaché *et al*. 2021), penguins (Mays *et al*. 2019), and flying lizards (Reilly *et al*. 2022). Furthermore, there appears to be significant variation in *gdi* within clades and between them. For example, in a survey of bird species the median *gdi* was 0.346 (1st quartile: 0.012; 3rd quartile: 0.742), while the surveyed mammalian species had a median *gdi* value of 0.799 (1st quartile: 0.716; 3rd quartile: 0.955) (Jackson *et al*. 2017). This suggests that appropriate heuristic cutoffs for *gdi* may differ among taxa, given that divergence times, population sizes and life histories differ substantially across these clades. Likewise, island radiations may also have a different underlying distribution of *gdi* given their shallow divergence times and propensity for rapid adaptation. Before a *gdi* heuristic can be confidently applied to Galapagos giant tortoise species, more thorough surveys of *gdi* among testudine species and across island radiations are needed.

When sister taxa are not species, Leaché *et al*. (2019) found that collapsing the taxa and recalculating *gdi* results in a larger *gdi*, signaling better support for the resulting species. This behavior of *gdi* can therefore be used to interpret the *gdi* without relying as heavily on heuristic thresholds. When we replicate this iterative process in BPP with our taxon set, we find that this pattern occurs when we collapse all central and southern Isabela Island taxa (i.e., *guntheri*, *microphyes*, *vandenburghi*, *vicina*) and when we collapse the Santa Cruz taxa (*porteri* and *donfaustoi*). All other collapses, however, result in estimates of *gdi* that are smaller rather than larger than the *gdi* of one or both sister taxa (Figure 3).

Applying population genetic principles to the *gdi* equation reveals why this occurs. Because *gdi* is calculated from the ratio of *τ* to *θ*, it will remain constant when these values change proportionally (e.g., the *gdi* will be 0.33 when *τ* is 0.001 and *θ* is 0.005, and when *τ* is 0.002 and *θ* is 0.01). When sister taxa are collapsed, both *τ* and *θ* will increase. By definition, in recent radiations *τ* will increase only marginally at each successive node. On the other hand, *θ* will be inflated when calculated from a highly structured sample (e.g., differentiated species) because the number of segregating sites is expected to be higher in a combined sample. Thus, the lower *gdi* values that we found for successive collapsing of taxa from different islands further supports the idea that these taxa do not belong to a single species.

The analyses presented in this study represent the most thorough attempt to date to tackle the question of species delimitation in Galapagos giant tortoises using genetic data. Several prior attempts to delimit Galapagos giant tortoise species focused on population genetic clustering, mitochondrial monophyly, and amount of sequence divergence (e.g., Caccone *et al*. 1999, Poulakakis *et al*. 2015, Loire *et al*. 2013, Kehlmaier *et al*. 2021). While these types of information can provide some evidence concerning species boundaries, they lack a strong foundation in modern population genetic theory as applied to species delimitation.

One such study claimed that genomic differentiation did not exist among *becki*, *vandenburghi*, and *porteri* (Loire *et al*. 2013), but a re-analysis of the data with appropriate filtering showed clear genomic differentiation of the three taxa (see Supplemental Figure S6 in Gaughran *et al*. 2018). Another study comparing only mitochondrial DNA data between Galapagos giant tortoises and extinct Caribbean giant tortoises argued that the recent divergence times of Galapagos giant tortoise taxa relative to other tortoise species disqualified them from species status (Kehlmaier *et al*. 2021). This view of species delimitation, however, ignores the complex reality of speciation (see Donoghue 1985, Mallet 1995, Hey 2006). In addition, it ignores the fact that some clades, such as those that colonized islands or other new environments, may be subject to different demographic and selective forces that are associated with adaptive radiations. Outside of tortoises, such a view would require a drastic re-delimitation of recently radiated species, with a disproportionate effect on small, endangered island populations.

To date, all published genetic evidence, including the data from Loire *et al*. (2013), supports the idea that Galapagos tortoise taxa are genetically distinct populations. Given the statistical power of genome-wide SNPs to accurately detect differentiation (Gaughran *et al*. 2018), this pattern of genetic distinctiveness appears unlikely to be overturned. Moving forward, genomic discussions of Galapagos giant tortoise taxonomy should recognize that the preponderance of evidence supports several genetically distinct taxa. However, whether these genetically distinct populations deserve to be considered species is a more nuanced question.

The unified species concept (de Queiroz 2005b, 2007) provides one way to understand this question. It highlights the idea of an independently evolving metapopulation lineage as the keystone property of every species concept, with other properties (e.g., morphological distinctiveness, genetic divergence, reproductive isolation) representing lines of evidence that a taxon is an independently evolving lineage. The unified species concept thereby provides a conceptual framework for evaluating candidate species that are morphologically similar, recently diverged, or continuing to hybridize. The evidence we present here, combined with decades of work documenting the distinctiveness of these taxa across multiple axes, shows how the Galapagos giant tortoise taxa meet many criteria discussed as important aspects of modern species concepts.

As described above, the diverse methods we apply here are not magic solutions to species delimitation, and each is the subject of ongoing debate. Still, interpreting our results holistically provides some clarification on species delimitation in Galapagos giant tortoises and highlights some areas of the taxonomy that remain difficult to resolve with our current data. Importantly, our results largely refute the single-species model of Galapagos giant tortoise taxonomy that was recently adopted by the IUCN Turtle Taxonomy Working Group (Rhodin *et al*. 2021). Instead, our modeling in BPP delimits at least 12 species and our modeling in PHRAPL delimits 13 species. On the other hand, the *gdi* results from BPP suggest 9 species, while those from PHRAPL suggest between 7 and 9 species (Figure 2). We find significant support for the species status of some taxa, especially the cases in which there is a single taxon per island. On the other hand, we find mixed support for taxa inhabiting the same island as different species (i.e., Isabela Island and Santa Cruz Island). Although model selection in BPP and PHRAPL supports the delimitation of species within islands, the pattern of *gdi* results suggests that Santa Cruz Island may be home to a single species and that the taxa of central and southern Isabela Island may also be a single species. Future work, incorporating more samples for each taxon, will likely resolve the ambiguous delimitations by making clear if the populations on Santa Cruz Island and central/southern Isabela Island have been most affected by constant migration, secondary contact after divergence, or other demographic processes.

Fundamentally, the taxonomic designation of Galapagos giant tortoises is both a scientific and a philosophical question, and one that is deserving of genuine debate in the literature. Still, this specific debate must be informed by decades of broader debates on species definitions and methods of species delimitation. The rich literature around this topic highlights the fact that speciation is a complex process: it can proceed at different rates and under different circumstances, and our own temporally-biased observations mean that we necessarily study taxa at different stages of this process. Galapagos giant tortoise taxa are evidently at different stages of lineage separation and divergence and therefore offer an exciting system in which to study both species boundaries and the process of speciation.

## Acknowledgements

We thank the Galápagos National Park Directorate for their collaboration. This work was supported by the Galapagos Conservancy (AWD0008098), RE:wild (AWD0008746), the Turtle Conservancy, and the Oak Foundation. SJG was supported by the National Science Foundation under Grant No. 2010918. We are grateful for input from Adam Leaché, Tomas Flouri, Ariadna Morales, and Bridgett vonHoldt in the development of this manuscript. This research made use of the Rocket High Performance Computing service at Newcastle University, the Yale Center for Research Computing, and Princeton Research Computing.

## Author Contributions

SJG, RG, AC, and ELJ designed the study. SJG, RG, AO, MJ, AC, and ELJ carried out the methodology. SJG, RG, MJ, NF, AO, JMM, NP, KdQ, and ELJ contributed to the investigation. SJG, RG, AO, MJ, NF, and ELJ visualized the results. SJG, AC, and ELJ acquired funding for the work. Project administration and supervision was provided by SJG, AC, and ELJ. SJG, RG, AO, MJ, AC, and ELJ wrote the original draft, and all authors contributed to reviewing and editing subsequent drafts.

## Data Accessibility

The whole genome resequencing data from Jensen *et al*. (2022) is available under NCBI BioProject PRJNA761229. Code for running BPP and PHRAPL, as well as alignments for loci used in these analyses, are available at: https://github.com/sjgaughran/tortoise-species-delimitation. Output files from SNAPP and BPP are available at: https://doi.org/10.5061/dryad.63xsj3v84

## Supplemental Methods

### Terminology, study design and data

We use several terms to refer to the demographic and evolutionary units of tortoises living in the Galapagos. We use the term *population* in the evolutionary genetics sense of a group of interbreeding individuals existing together in time and space (e.g., Hedrick 2009). We use the term *lineage* in a slightly broader sense, to expand the time component of *population*, thereby including ancestor-descendant relationships. We use the systematic term *taxon* to refer broadly to an evolutionary unit (e.g., lineage) without specifying whether that unit is a species or a subspecies. Because our goal is to evaluate the status of Galapagos giant tortoise taxa as species, we avoid using the terms *species* and *subspecies* as *a priori* descriptors. To maintain this agnosticism, we use only the epithet to refer to each taxon (e.g., *phantasticus* rather than *Chelonoidis phantasticus* or *Chelonoidis niger phantasticus*).

We designed a whole-genome study of population distinctiveness and species delimitation, taking advantage of existing illumina short-read whole genome resequencing data (Jensen *et al*. 2021, Jensen *et al*. 2022), which are available under NCBI Bioproject PRJNA761229. The samples in our data set represent all extant taxa, which have previously been classified as populations, species, or subspecies. The samples were originally selected for sequencing as representatives of the genetic and geographic populations (Figure 1) that have previously been shown to exist using microsatellite (Ciofi *et al*. 2002, Ciofi *et al*. 2006) and reduced representation SNP (Miller *et al*. 2018) data. Previous genetic research suggests that the *becki* taxon consists of two geographically and genetically distinct populations, referred to in the literature as “PBL” (Piedras Blancas) and “PBR” (Puerto Bravo) (Garrick *et al*. 2014). A prior phylogeny constructed from these genome data in Jensen *et al*. (2022) showed the named taxa to be monophyletic, with the exception of *becki*. Within *becki*, PBR individuals formed a clade sister to *darwini* and PBL individuals did not form a clade, with one sample closest to PBR and *darwini* and the other two closest to other taxa from Isabela. We have therefore chosen to test *becki (PBL)* and *becki (PBR)* as potentially distinct taxa, and have retained the potentially admixed PBL individual to avoid biasing our analyses.

We mapped these reads to a reference genome for *abingdonii* (Quesada et al. 2019) using bwa-mem (Li 2013). Jensen *et al*. (2021) previously investigated the potential for reference bias in using the *abingdonii* assembly and found no evidence for such a bias. We called SNPs using the BCFtools variant calling pipeline (Li et al. 2009). We then filtered the VCF for bi-allelic SNPs that had a minimum GQ of 25, a minimum map quality score of 25, and at least 2 reads for each allele in the genotype. We then created a consensus fasta file for each individual using BCFtools and masked each consensus sequence for all missing genotypes from the VCF to ensure that missing data were not erroneously assigned to the reference allele. We also masked every fasta with a mask of repetitive regions downloaded from the UCSC Genome Browser and with a mappable regions mask generated by Jensen *et al*. (2021). From these filtered and masked fastas, we generated a random set of 1kb loci separated by 100kb. We retained loci that had less than 10% missing data, GC content between 30% and 70%, and at least one variable site. Because BPP requires phased data when using more than ∼100 loci, we phased each locus using PHASE v. 2.1.1 (Stephens *et al*. 2001, Stephens and Scheet 2005). We then used several methods to delimit species under different phylogenetic and coalescent models.

### SNAPP Tree Generation

We used SNAPP v1.5.2 (Bryant *et al*. 2012) and MODEL_SELECTION v1.5.3 (Baele *et al*. 2012), as implemented in BEAST2 v2.6.7 (Bouckaert *et al*. 2014), to generate a guide tree reflecting the phylogenetic relationships among the extant populations of Galapagos giant tortoises for downstream species delimitation comparisons. We first tested 21 different phylogenetic models on a dataset comprising 1,000 random, unlinked SNPs—and corresponding genotypes— from 39 samples. We obtained this dataset from a previous study conducted by Jensen et al. (2022) and it included one Chaco tortoise (*C. chilensis*) sample to serve as an outgroup to the Galapagos giant tortoises. We ensured the Galapagos giant tortoises presented genetic variation across loci and we allowed no missing data.

We grouped the samples into populations, assumed forward and backward mutation rates equivalent to the unit (*u* = *v* = 1), and used combinations of shape (α = 5, 12, 30), scale (β = 50, 60, 70, 80, 110, 150), and speciation rate (λ = 0.01, 10, 39) parameters as priors to constitute the models to be tested (see Table S1). Path sampling runs for each model consisted of 24 steps, 100,000 MCMC generations sampled every 100 generations, and a 50% burn-in. We compared the resulting marginal likelihood estimates from each run and selected the best model via Bayes factor delimitation (Kass & Raftery 1995) after using the default model (α = 12, β = 110, λ = 0.01) as a reference.

For the best model, we then used an additional set of 4,000 random, unlinked biallelic SNPs (5,000 SNPs total) for final phylogenetic reconstruction. For this dataset, we performed four independent runs consisting of 2,000,000 MCMC generations sampled every 1,000 generations and a 10% burn-in. We used Tracer v1.7.2 (Rambaut *et al*. 2018) to assess statistical convergence across runs and LogCombiner v2.6.7 (Drummond & Rambaut 2007) to summarize the posterior trees.

### Divergence time from BPP

We can calculate the absolute divergence time in generations from τ, the rate-scaled divergence time in BPP, as:

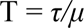

Where *µ* is the per-generation mutation rate. Although the per-generation mutation rate in Galapagos giant tortoises is unknown, we use 6×10^−9^ reflecting the *de novo* mutation rate recently measured in the painted turtle (Bergeron *et al*. 2023).

## Supplemental Figures and Tables

**Figure S1.**
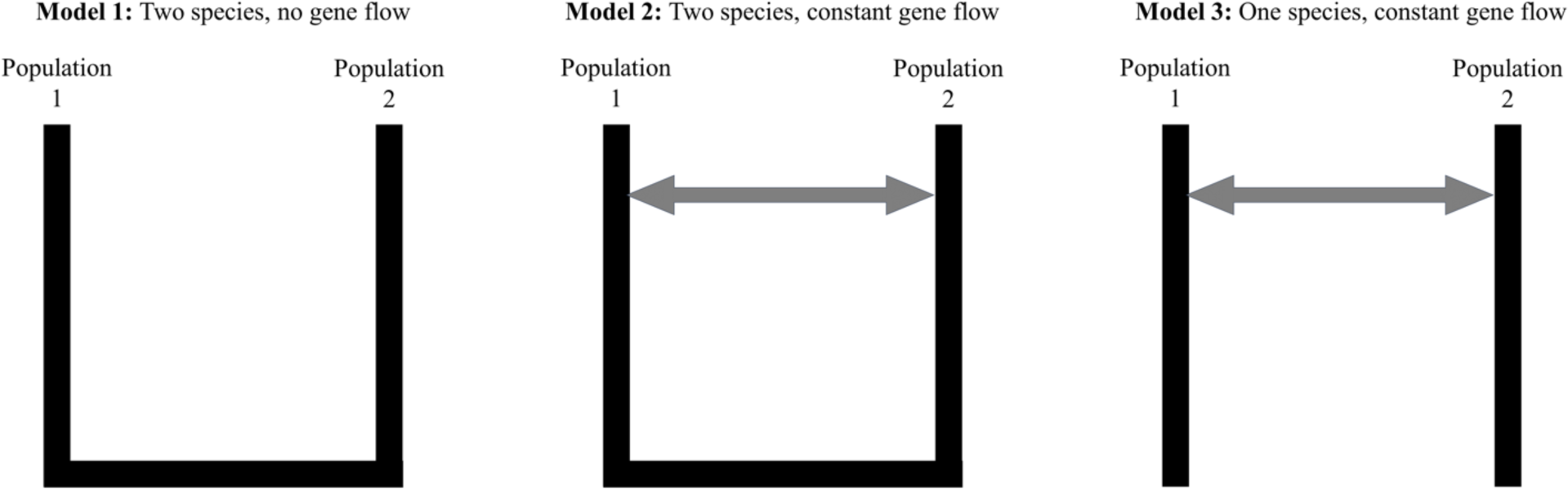
The three models used to test one and two species models under differing gene flow scenarios in PHRAPL. The grey arrow indicates bi-directional gene flow between the two populations.

**Figure S2.**
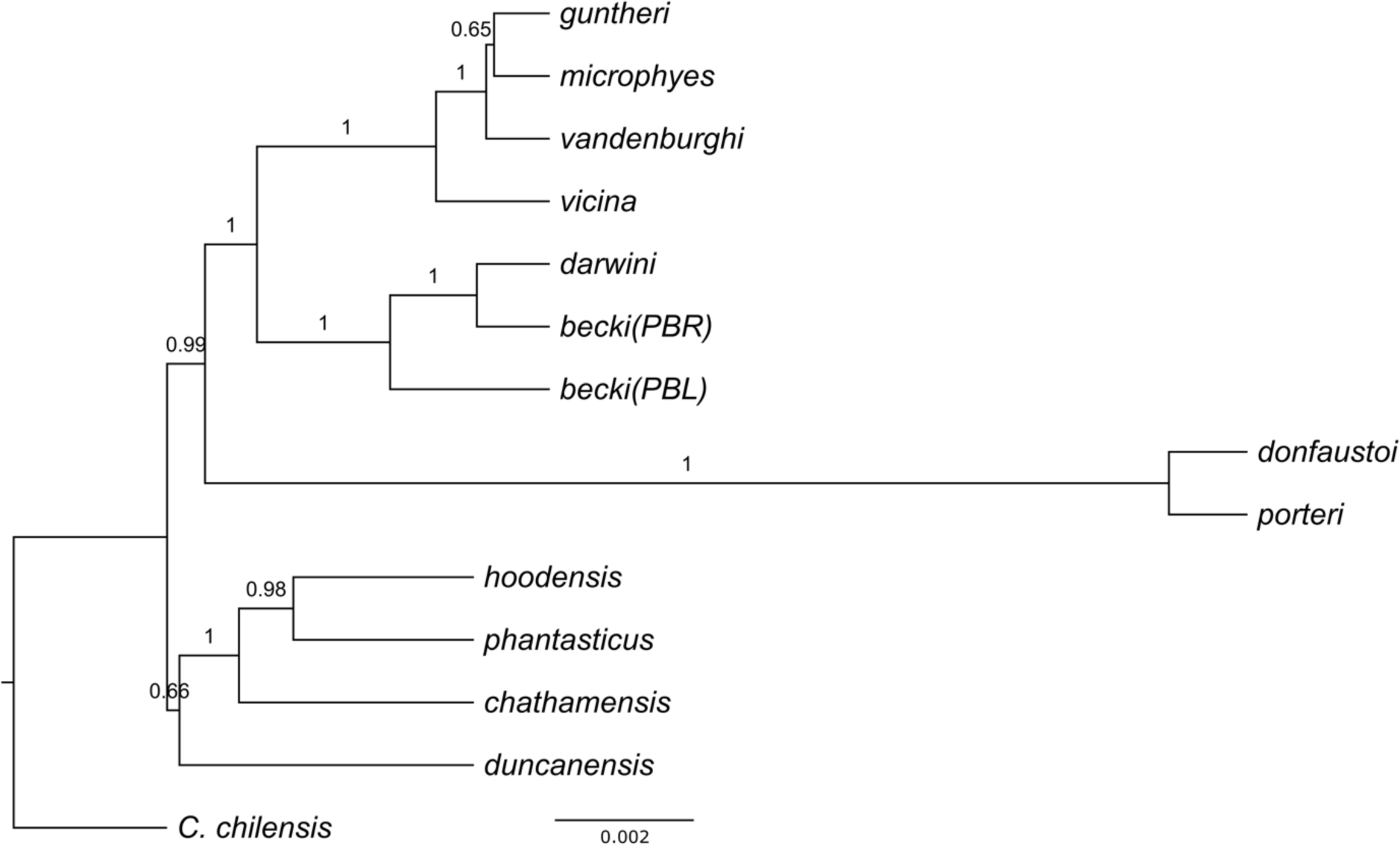
Phylogeny with scaled branch lengths generated from 5,000 genome-wide SNPs in SNAPP from the best supported model (α = 5, β = 150, λ = 39) with branch posterior support values indicated. The scale bar indicates the number of nucleotide substitutions per site.

**Figure S3.**
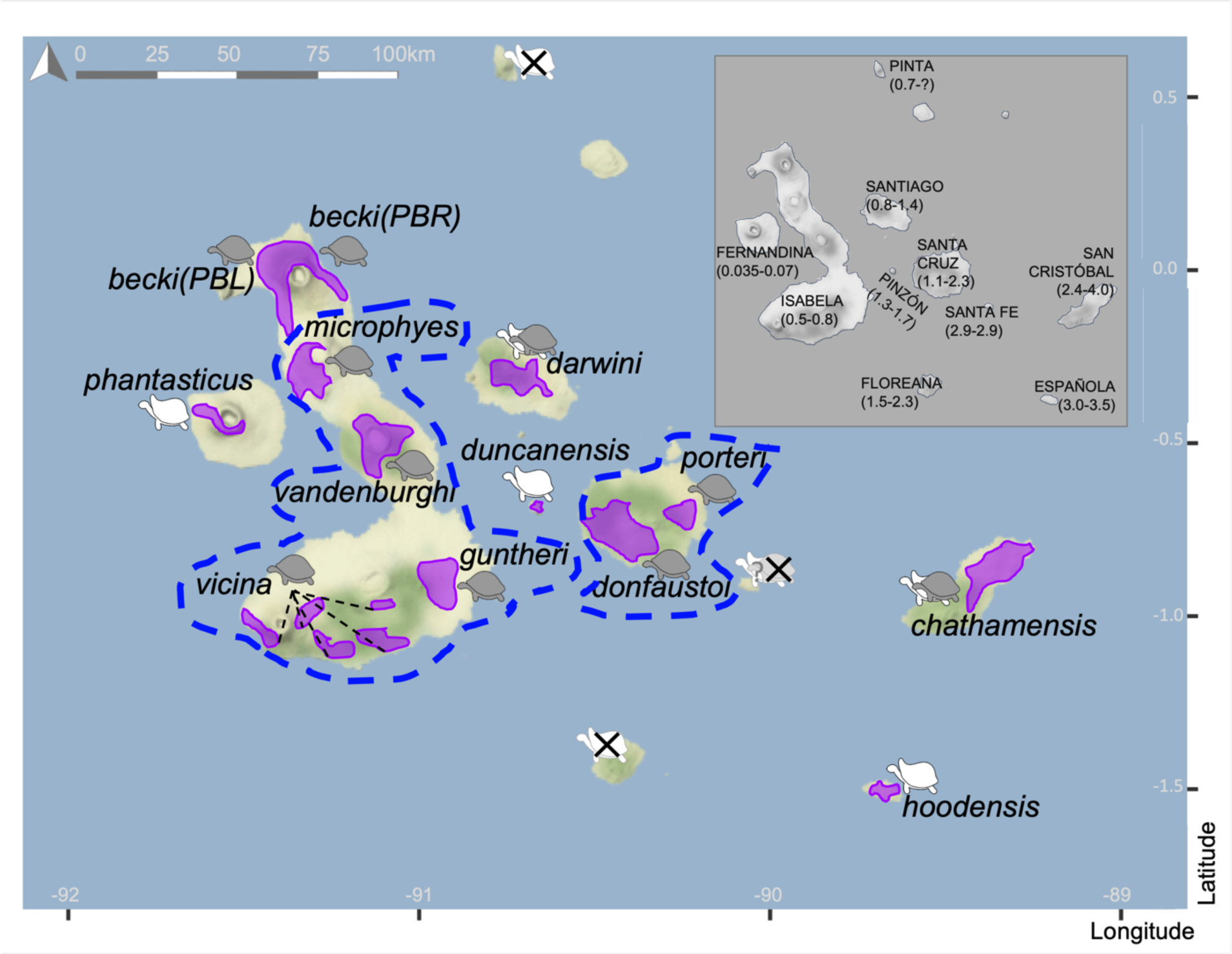
Galapagos giant tortoise species ranges when the Santa Cruz Island taxa (*porteri* and *donfaustoi*) are collapsed and the central and southern Isabela Island (*microphyes*, *vandenburghi*, *guntheri*, and *vicina*) are collapsed. Purple shapes indicate current ranges of populations. Blue dotted lines surround taxa that are collapsed into a single species according to some of our analyses. Carapace morphology (domed = gray, saddleback = white) is shown for each population, with “intermediate” shell shape indicated with overlapping icons. Icons with an “X” indicate extinct tortoise populations. Inset map shows the names of islands with recently extant Galapagos giant tortoise taxa, with geological ages of each island in million years below.

**Table S1.**
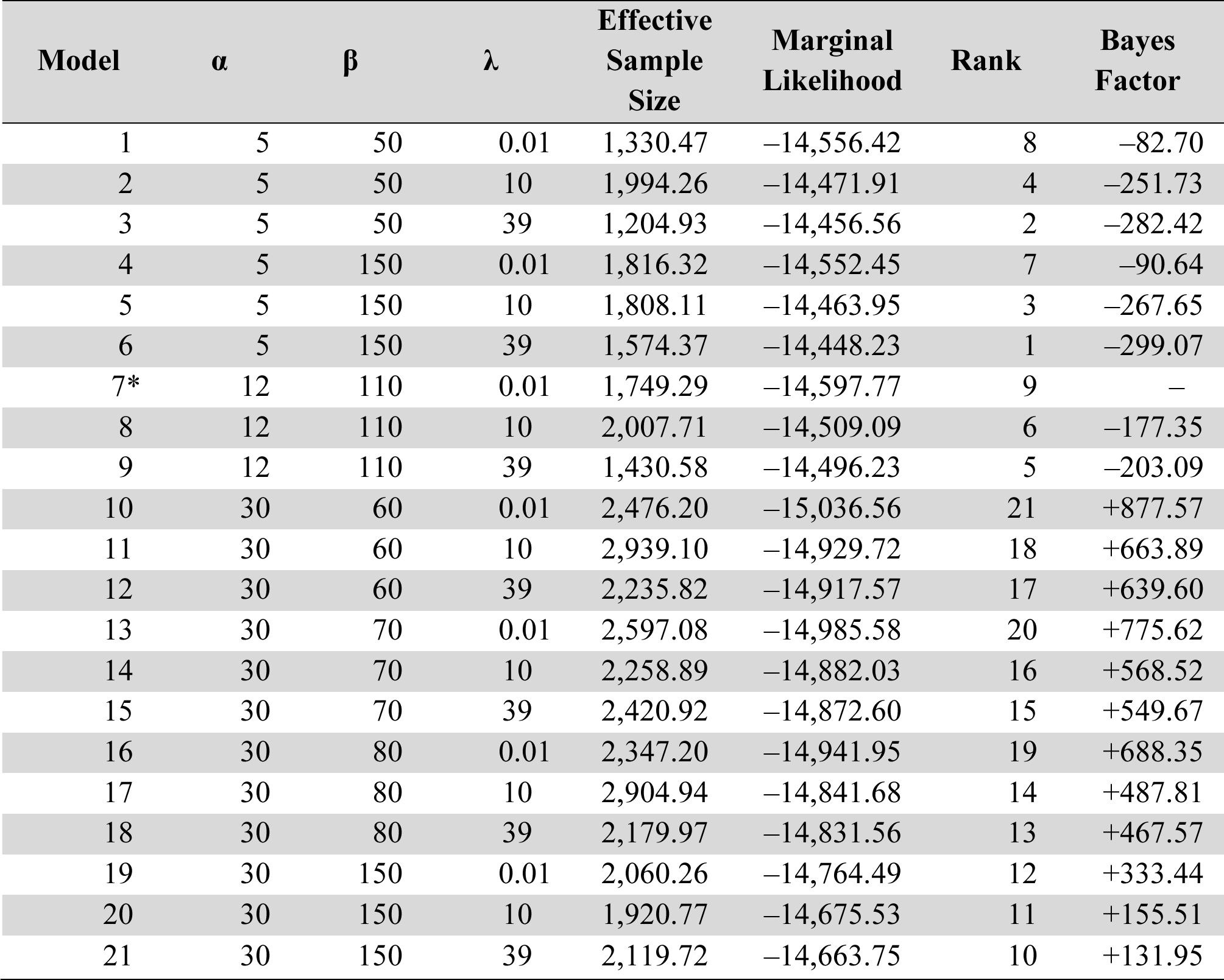
Summary statistics from 21 different phylogenetic models tested on 1,000 random, unlinked biallelic SNPs—and corresponding genotypes—from 39 tortoises arranged into populations. Each run consisted of 24 steps, 100,000 MCMC generations sampled every 100 generations, and a 50% burn-in. The default, reference model for Bayes factor delimitation is shown with an asterisk (*).

**Table S2.** Parameter values for migration (*m*) and divergence (*t*) used for the grid search within PHRAPL. The maximum divergence time is equivalent to 1Ma.

|  |  |  |  |  |  |  |  |
| --- | --- | --- | --- | --- | --- | --- | --- |
| m | 0.005 | 0.02 | 0.08 | 0.12 | 0.25 | 0.4 | 0.8 |
| t | 0.01 | 0.05 | 0.1 | 0.25 | 0.5 | 0.75 | 1 |

**Table S3.**
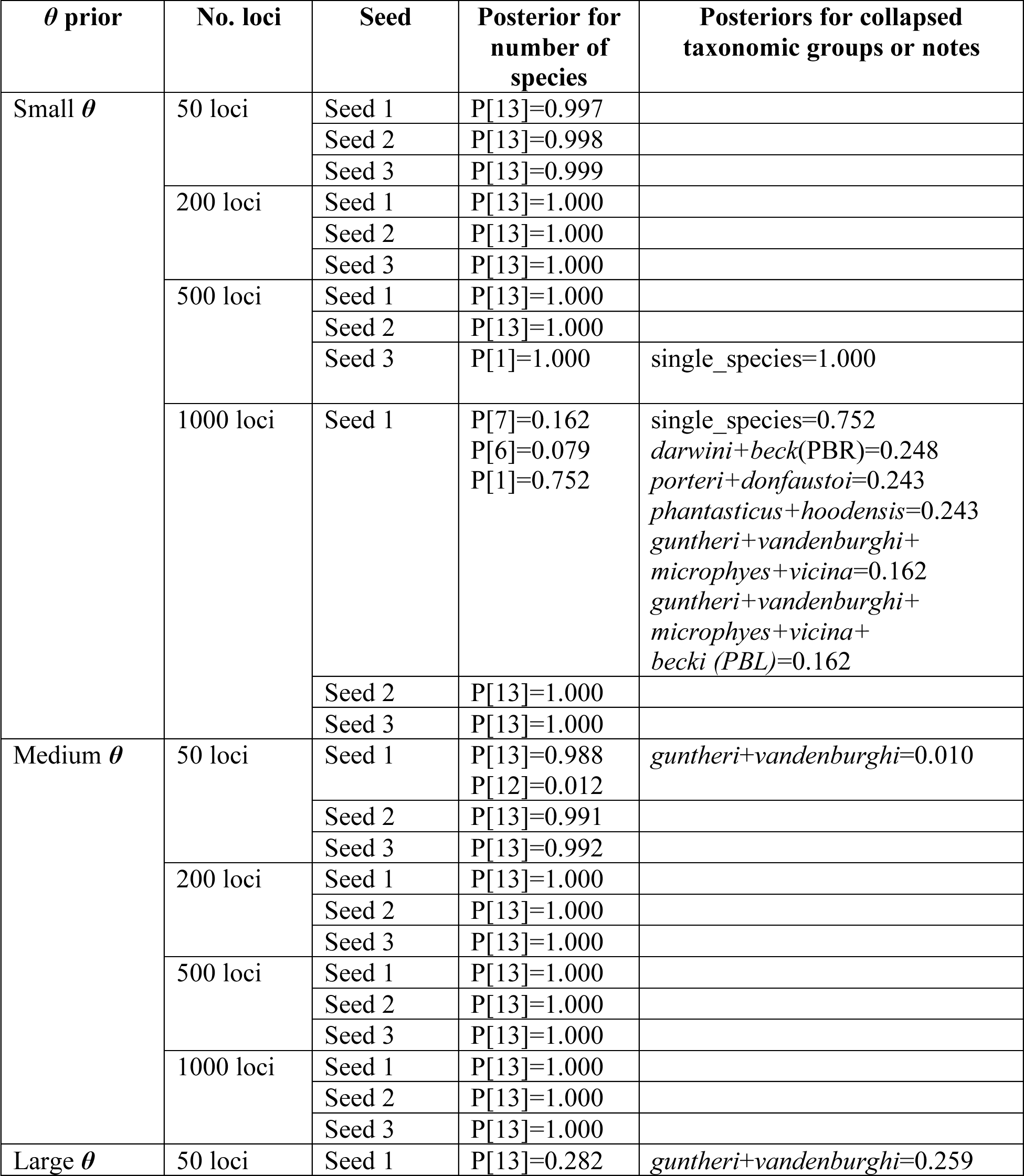
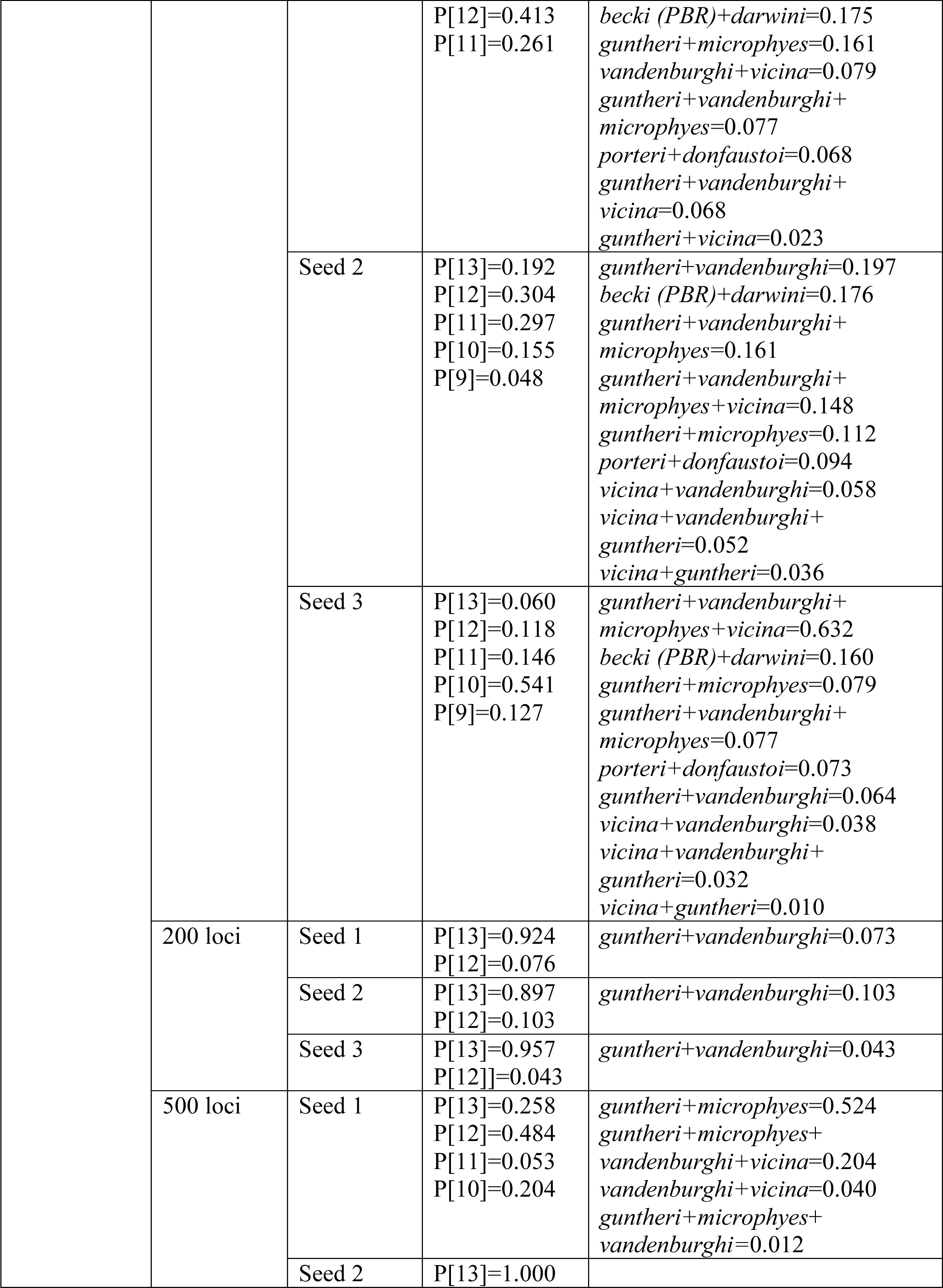
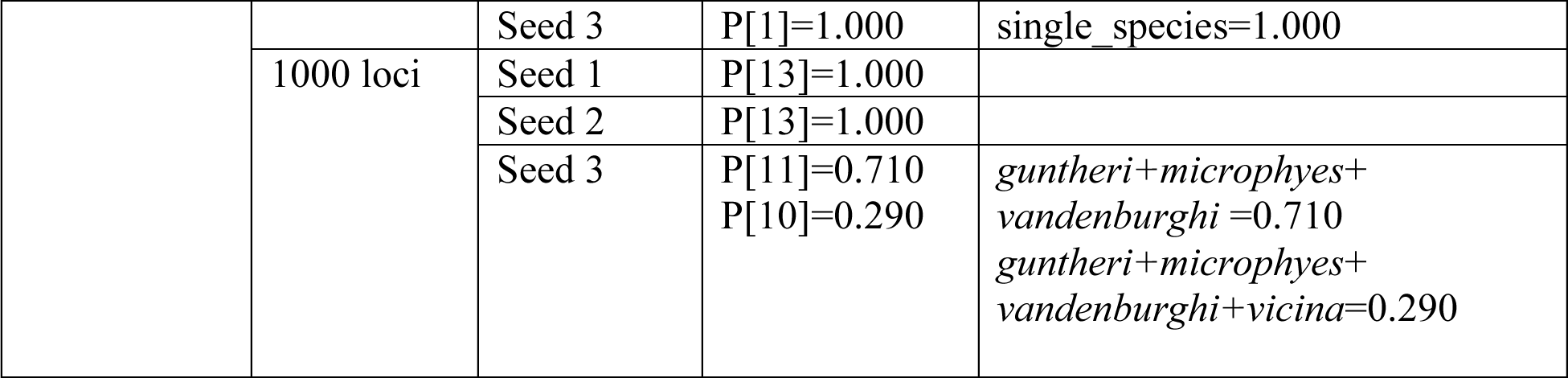
Posterior output from the A11 analysis species delimitation model in BPP using phased loci from the 38 Galapagos giant tortoise individuals. Support for collapsed taxa as species is included in the final column. The “Medium *θ*” prior is the most realistic prior.

**Table S4.**
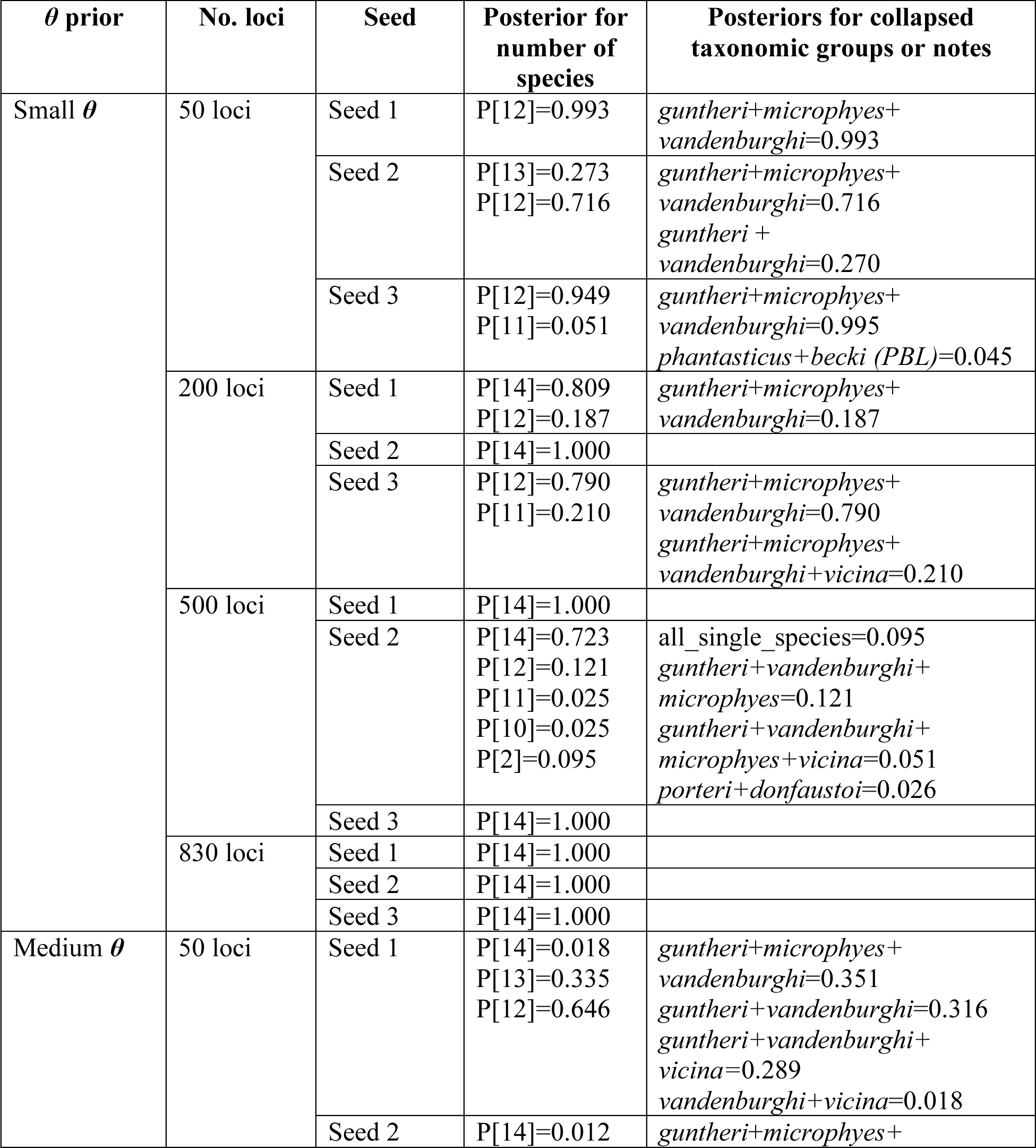
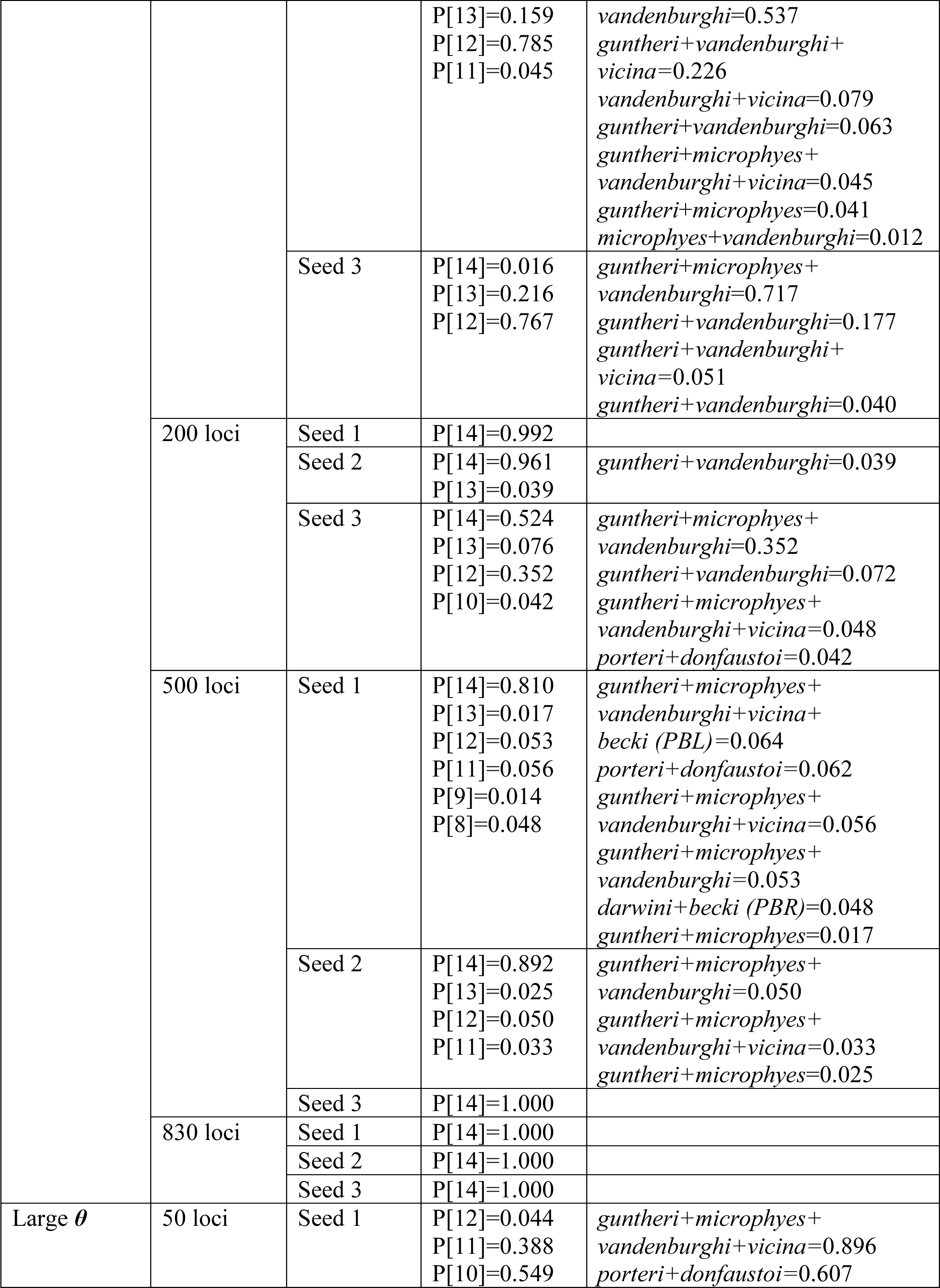
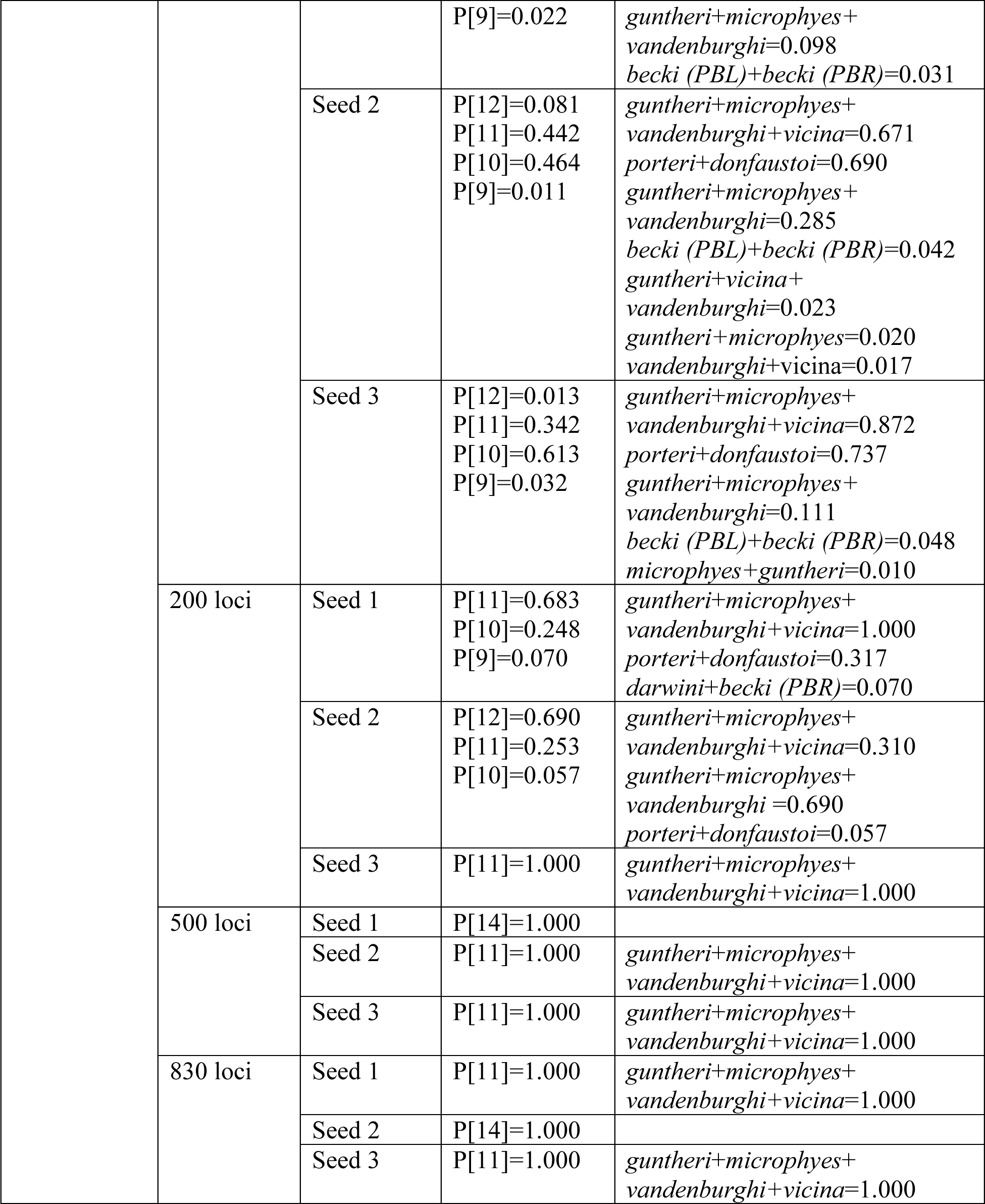
Posterior output from the A11 analysis species delimitation model in BPP using phased loci from the 38 Galapagos giant tortoise individuals and the Chaco tortoise individual. Support for collapsed taxa as species is included in the final column. The “Medium *θ*” prior is the most realistic prior.

**Table S5.** Posterior output from the A11 analysis species delimitation model in BPP using unphased loci from the 38 Galapagos giant tortoise individuals. Support for collapsed taxa as species is included in the final column. The “Medium *θ*” prior is the most realistic prior.

| $\theta$ prior | No. loci | Seed | Posterior for number of species | Posteriors for collapsed taxonomic groups or notes |
| --- | --- | --- | --- | --- |
| Small $\theta$ | 50 loci | Seed 1 | P[13]=0.986<br>P[12]=0.014 | *porteri*+*donfaustoi*=0.013 |
|  |  | Seed 2 | P[13]=0.997 |  |
|  |  | Seed 3 | P[13]=0.995 |  |
| Medium $\theta$ | 50 loci | Seed 1 | P[13]=0.992 | |
|  |  | Seed 2 | P[13]=0.993 |  |
|  |  | Seed 3 | P[1]=1.000 | single species=1.000 |
| Large $\theta$ | 50 loci | Seed 1 | P[13]=0.577<br>P[12]=0.357<br>P[11]=0.063 | *guntheri*+*vandenburghi*=0.228<br>*darwini*+*becki* (PBR)=0.056<br>*microphyes*+*guntheri*=0.037<br>*vandenburghi*+*vicina*=0.036<br>*guntheri*+*vicina*=0.036<br>*porteri*+*donfaustoi*=0.026<br>*guntheri*+*vandenburghi*+*vicina*=0.025 |
|  |  | Seed 2 | P[13]=0.517<br>P[12]=0.334<br>P[11]=0.066<br>P[10]=0.071<br>P[9]=0.012 | *guntheri*+*vandenburghi*=0.232<br>*darwini*+*becki* (PBR)=0.066<br>*guntheri*+*vandenburghi*+*microphyes*+*vicina*=0.075<br>*microphyes*+*guntheri*=0.050<br>*porteri*+*donfaustoi*=0.031<br>*guntheri*+*vicina*=0.030<br>*guntheri*+*vandenburghi*+*vicina*=0.029<br>*vandenburghi*+*vicina*=0.027 |
|  |  | Seed 3 | P[13]=0.493<br>P[12]=0.391<br>P[11]=0.106<br>P[10]=0.010 | *guntheri*+*vandenburghi*=0.248<br>*darwini*+*becki* (PBR)=0.092<br>*microphyes*+*guntheri*=0.064<br>*guntheri*+*vicina*=0.039<br>*guntheri*+*vandenburghi*+*vicina*=0.039<br>*vandenburghi*+*vicina*=0.036<br>*porteri*+*donfaustoi*=0.024<br>*guntheri*+*vandenburghi*+*microphyes* =0.022 |

**Table S6.**
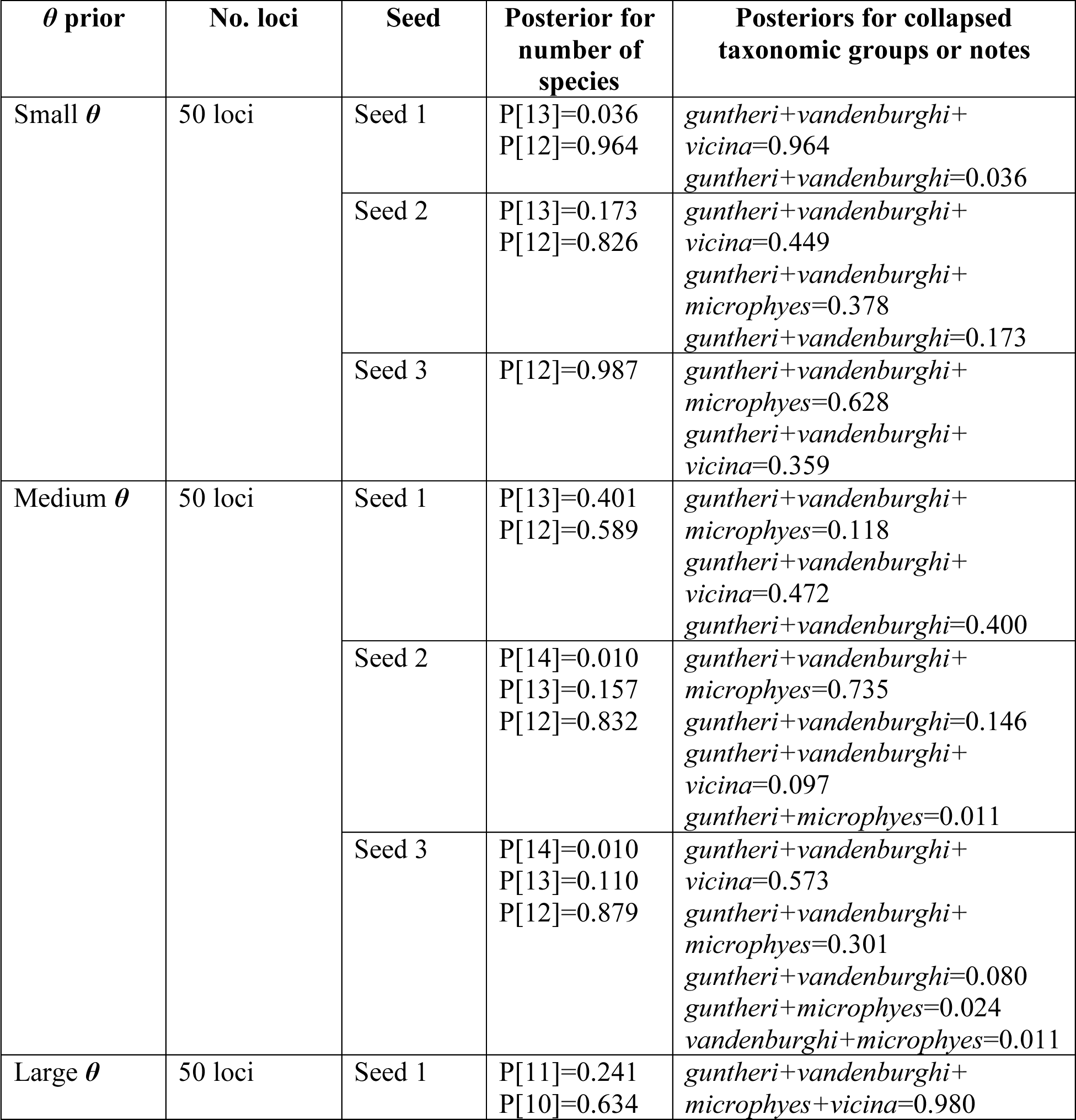
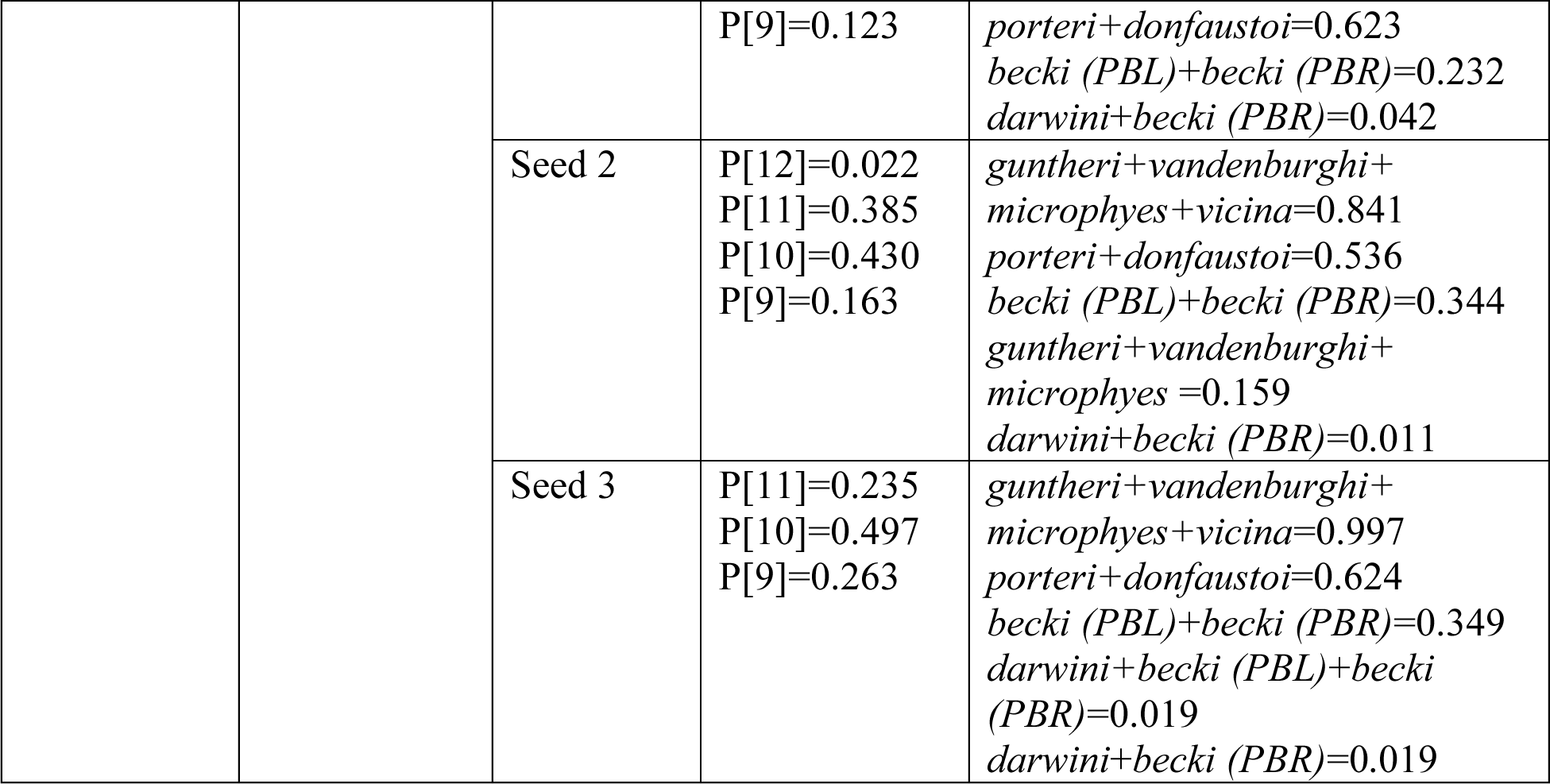
Posterior output from the A11 analysis species delimitation model in BPP using unphased loci from the 38 Galapagos giant tortoise individuals and the Chaco tortoise individual. Support for collapsed taxa as species is included in the final column. The “Medium *θ*” prior is the most realistic prior.

**Table S7.** Posterior output from the A10 analysis species delimitation model in BPP using phased loci from the 38 Galapagos giant tortoise individuals. Nodes in the guide tree with a posterior probability <0.95 were counted as a single species. The “Medium *θ*” prior is the most realistic prior. A maximum of 13 species (i.e., each Galapagos giant tortoise taxon) was possible.

| $\theta$ prior | No. loci | Seed | Number of species<br>with >0.95<br>posterior support |
| --- | --- | --- | --- |
| Small $\theta$ | 50 loci | Seed 1 | 13 |
|  |  | Seed 2 | 13 |
|  |  | Seed 3 | 13 |
|  | 200 loci | Seed 1 | 13 |
|  |  | Seed 2 | 13 |
|  |  | Seed 3 | 13 |
|  | 500 loci | Seed 1 | 13 |
|  |  | Seed 2 | 13 |
|  |  | Seed 3 | 13 |
|  | 1000 loci | Seed 1 | 13 |
|  |  | Seed 2 | 13 |
|  |  | Seed 3 | 13 |
| Medium $\theta$ | 50 loci | Seed 1 | 13 |
|  |  | Seed 2 | 13 |
|  |  | Seed 3 | 13 |
|  | 200 loci | Seed 1 | 13 |
|  |  | Seed 2 | 13 |
|  |  | Seed 3 | 13 |
|  | 500 loci | Seed 1 | 13 |
|  |  | Seed 2 | 13 |
|  |  | Seed 3 | 13 |
|  | 1000 loci | Seed 1 | 13 |
|  |  | Seed 2 | 13 |
|  |  | Seed 3 | 13 |
| Large $\theta$ | 50 loci | Seed 1 | 8 |
|  |  | Seed 2 | 9 |
|  |  | Seed 3 | 9 |
|  | 200 loci | Seed 1 | 11 |
|  |  | Seed 2 | 10 |
|  |  | Seed 3 | 11 |
|  | 500 loci | Seed 1 | 12 |
|  |  | Seed 2 | 12 |
|  |  | Seed 3 | 12 |
|  | 1000 loci | Seed 1 | 12 |
|  |  | Seed 2 | 13 |
|  |  | Seed 3 | 13 |

**Table S8.** Posterior output from the A10 analysis species delimitation model in BPP using phased loci from the 38 Galapagos giant tortoise individuals and the Chaco tortoise individual. Nodes in the guide tree with a posterior probability <0.95 were counted as a single species. The “Medium *θ*” prior is the most realistic prior. A maximum of 14 species (i.e., each Galapagos giant tortoise taxon and the Chaco tortoise) was possible.

| $\theta$ prior | No. loci | Seed | Number of species<br>with >0.95<br>posterior support |
| --- | --- | --- | --- |
| Small $\theta$ | 50 loci | Seed 1 | 10 |
|  |  | Seed 2 | 11 |
|  |  | Seed 3 | 12 |
|  | 200 loci | Seed 1 | 14 |
|  |  | Seed 2 | 13 |
|  |  | Seed 3 | 11 |
|  | 500 loci | Seed 1 | 10 |
|  |  | Seed 2 | 12 |
|  |  | Seed 3 | 10 |
|  | 830 loci | Seed 1 | 14 |
|  |  | Seed 2 | 10 |
|  |  | Seed 3 | 14 |
| Medium $\theta$ | 50 loci | Seed 1 | 12 |
|  |  | Seed 2 | 12 |
|  |  | Seed 3 | 12 |
|  | 200 loci | Seed 1 | 14 |
|  |  | Seed 2 | 14 |
|  |  | Seed 3 | 12 |
|  | 500 loci | Seed 1 | 14 |
|  |  | Seed 2 | 14 |
|  |  | Seed 3 | 14 |
|  | 830 loci | Seed 1 | 14 |
|  |  | Seed 2 | 14 |
|  |  | Seed 3 | 14 |
| Large $\theta$ | 50 loci | Seed 1 | 9 |
|  |  | Seed 2 | 10 |
|  |  | Seed 3 | 10 |
|  | 200 loci | Seed 1 | 12 |
|  |  | Seed 2 | 12 |
|  |  | Seed 3 | 11 |
|  | 500 loci | Seed 1 | 11 |
|  |  | Seed 2 | 13 |
|  |  | Seed 3 | 12 |
|  | 830 loci | Seed 1 | 13 |
|  |  | Seed 2 | 11 |
|  |  | Seed 3 | 12 |

**Table S9.** Divergence time estimates in generations from BPP, with prior *θ* ∼ IG(3, 0.001) and prior *τ* ∼ IG(3, 0.001). Median and 95% credible interval calculated from the posterior distribution of *τ*. Divergence time calculated using a per-generation mutation rate of 6×10^−9^.

| **Taxon** | **Median divergence time in generations**<br>**(95% credible interval)** |
| --- | --- |
| *darwini* | 2667 (2000–3333) |
| *becki (PBR)* | 2667 (2000–3333) |
| *guntheri* | 1333 (1167–1667) |
| *microphyes* | 1333 (1167–1667) |
| *vandenburghi* | 1500 (1333–1667) |
| *vicina* | 2333 (2000–2667) |
| *becki (PBL)* | 3833 (3167–4667) |
| *donfaustoi* | 3167 (2500–4000) |
| *porteri* | 3167 (2500–4000) |
| *hoodensis* | 4500 (3500–5500) |
| *phantasticus* | 4500 (3500–5500) |
| *duncanensis* | 5667 (4833–6833) |
| *chathamensis* | 6333 (5500–7833) |
| *C. chilensis* | 368,500 (352,667–392,167) |

**Table S10.** Effective population size estimates from BPP, with prior *θ* ∼ IG(3, 0.001) and prior *τ* ∼ IG(3, 0.001) and an assumed mutation rate of 6.0 x 10^−9^. Median and 95% credible interval calculated from the posterior distribution of *θ*. Effective population size calculated using a per-generation mutation rate of 6×10^−9^.

| **Taxon** | **Median effective population size**<br>**(95% credible interval)** |
| --- | --- |
| *darwini* | 8208 (6000–10,667) |
| *becki* (PBR) | 5458 (4083–6958) |
| *guntheri* | 6416 (4833–8833) |
| *microphyes* | 3917 (3125–5083) |
| *vandenburghi* | 12,125 (8583–19,042) |
| *vicina* | 27,333 (25,542–29,208) |
| *becki* (PBL) | 19,750 (15,208–25,875) |
| *donfaustoi* | 5083 (3958–6625) |
| *porteri* | 45,125 (26,083–94,333) |
| *hoodensis* | 2917 (2333–3542) |
| *phantasticus* | 94,042 (45,208–223,375) |
| *duncanensis* | 5750 (4875–7125) |
| *chathamensis* | 9333 (7833–11,375) |
| *C. chilensis* | 77,917 (71,292–85,292) |

**Table S11.** Estimates of *gdi* for each taxon, calculated from the posterior distributions of *θ* and *τ* in BPP, with prior *θ* ∼ IG(3, 0.001) and prior *τ* ∼ IG(3, 0.001).

| **Taxon** | **Median *gdi***<br>**(95% credible interval)** |
| --- | --- |
| *darwini* | 0.149 (0.140–0.159) |
| *becki (PBR)* | 0.218 (0.187–0.247) |
| *guntheri* | 0.104 (0.077–0.131) |
| *microphyes* | 0.162 (0.132–0.194) |
| *vandenburghi* | 0.063 (0.040–0.087) |
| *vicina* | 0.107 (0.082–0.134) |
| *becki (PBL)* | 0.094 (0.075–0.115) |
| *donfaustoi* | 0.266 (0.231–0.278) |
| *porteri* | 0.034 (0.016–0.058) |
| *hoodensis* | 0.534 (0.497–0.572) |
| *phantasticus* | 0.023 (0.010–0.049) |
| *duncanensis* | 0.423 (0.393–0.455) |
| *chathamensis* | 0.262 (0.236–0.290) |

**Table S12.**
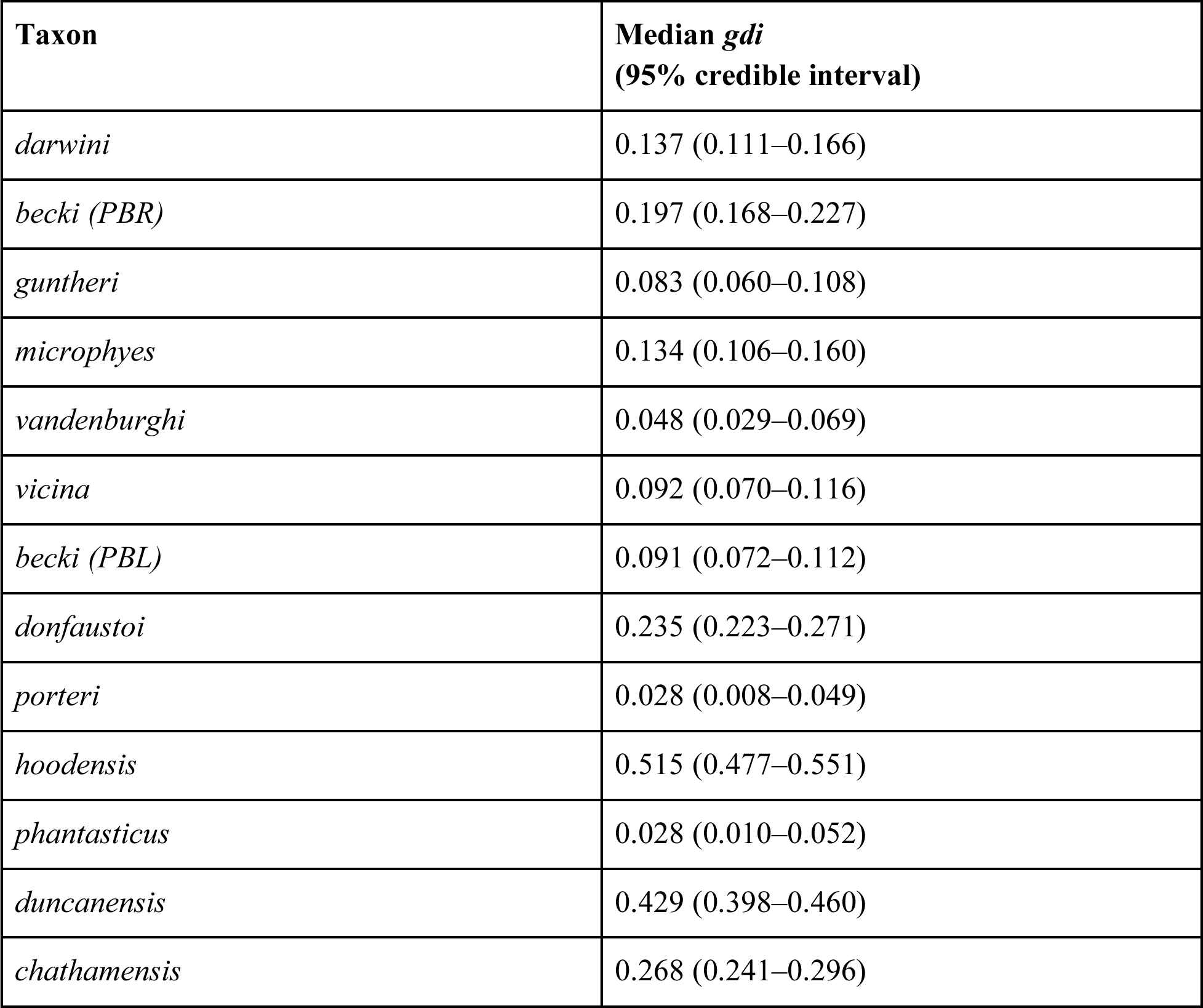
Estimates of *gdi* for each taxon, calculated from the posterior distributions of *θ* and *τ* in BPP, with prior *θ* ∼ IG(3, 0.005) and prior *τ* ∼ IG(3, 0.001).

**Table S13.** Estimates of *gdi* for each taxon, calculated from the posterior distributions of *θ* and *τ* in BPP, with prior *θ* ∼ IG(3, 0.0001) and prior *τ* ∼ IG(3, 0.001).

| **Taxon** | **Median *gdi***<br>**(95% credible interval)** |
| --- | --- |
| *darwini* | 0.152 (0.124–0.180) |
| *becki* (PBR) | 0.223 (0.193–0.255) |
| *guntheri* | 0.109 (0.077–0.143) |
| *microphyes* | 0.171 (0.133–0.210) |
| *vandenburghi* | 0.064 (0.041–0.090) |
| *vicina* | 0.107 (0.081–0.135) |
| *becki* (PBL) | 0.094 (0.074–0.115) |
| *donfaustoi* | 0.271 (0.233–0.309) |
| *porteri* | 0.032 (0.012–0.058) |
| *hoodensis* | 0.537 (0.502–0.573) |
| *phantasticus* | 0.025 (0.013–0.046) |
| *duncanensis* | 0.413 (0.381–0.446) |
| *chathamensis* | 0.256 (0.228–0.283) |

**Table S14.** Estimates of *gdi* for each taxon after the first round of sister taxa were collapsed. The *gdi* was calculated from the posterior distributions of *θ* and *τ* of the collapsed taxa in BPP, with prior *θ* ∼ IG(3, 0.001) and prior *τ* ∼ IG(3, 0.001).

| **Taxon** | **Median *gdi***<br>**(95% credible interval)** |
| --- | --- |
| *darwini+becki (PBR)* | 0.161 (0.144–0.180) |
| *guntheri+microphyes* | 0.044 (0.029–0.062) |
| *vandenburghi* | 0.158 (0.127–0.190) |
| *vicina* | 0.105 (0.080–0.131) |
| *becki (PBL)* | 0.093 (0.074–0.113) |
| *donfaustoi+porteri* | 0.291 (0.268–0.314) |
| *hoodensis+phantasticus* | 0.190 (0.170–0.211) |
| *duncanensis* | 0.427 (0.396–0.460) |
| *chathamensis* | 0.263 (0.235–0.291) |

**Table S15.**
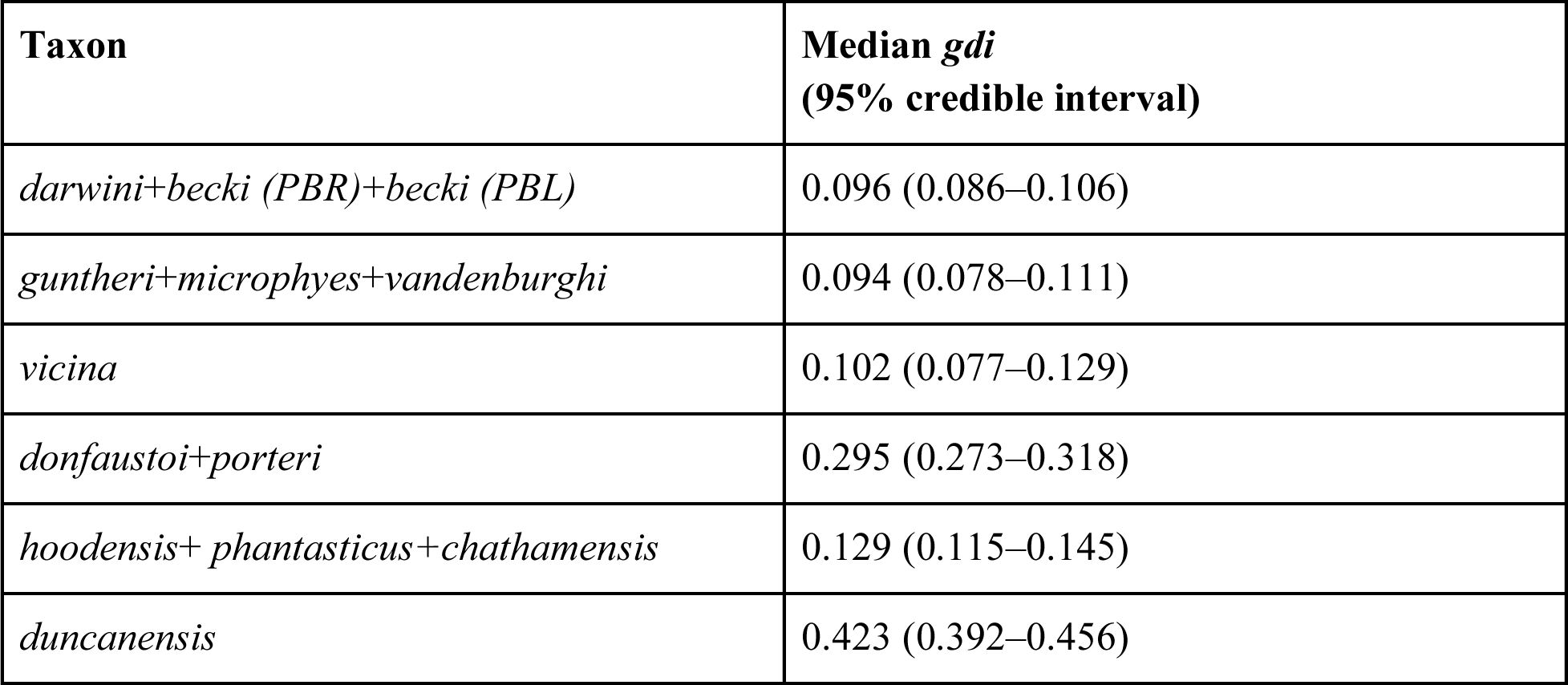
Estimates of *gdi* for each taxon after the second round of sister taxa were collapsed. The *gdi* was calculated from the posterior distributions of *θ* and *τ* of the collapsed taxa in BPP, with prior *θ* ∼ IG(3, 0.001) and prior *τ* ∼ IG(3, 0.001).

| **Taxon** | **Median *gdi***<br>**(95% credible interval)** |
| --- | --- |
| *darwini+becki (PBR)+becki (PBL)* | 0.096 (0.086–0.106) |
| *guntheri+microphyes+vandenburghi* | 0.094 (0.078–0.111) |
| *vicina* | 0.102 (0.077–0.129) |
| *donfaustoi+porteri* | 0.295 (0.273–0.318) |
| *hoodensis+ phantasticus+chathamensis* | 0.129 (0.115–0.145) |
| *duncanensis* | 0.423 (0.392–0.456) |

**Table S16.** Estimates of *gdi* for each taxon after the third round of sister taxa were collapsed. The *gdi* was calculated from the posterior distributions of *θ* and *τ* of the collapsed taxa in BPP, with prior *θ* ∼ IG(3, 0.001) and prior *τ* ∼ IG(3, 0.001).

| **Taxon** | **Median *gdi***<br>**(95% credible interval)** |
| --- | --- |
| *darwini+becki (PBR)+becki (PBL)* | 0.099 (0.089–0.110) |
| *guntheri+microphyes+vandenburghi+vicina* | 0.226 (0.211–0.243) |
| *donfaustoi+porteri* | 0.298 (0.275–0.322) |
| *hoodensis+ phantasticus+duncanensis+ chathamensis* | 0.144 (0.134–0.156) |

**Table S17.** Estimates of *gdi* for each taxon after the fourth round of sister taxa were collapsed. The *gdi* was calculated from the posterior distributions of *θ* and *τ* of the collapsed taxa in BPP, with prior *θ* ∼ IG(3, 0.001) and prior *τ* ∼ IG(3, 0.001).

| **Taxon** | **Median *gdi***<br>**(95% credible interval)** |
| --- | --- |
| *darwini+becki (PBR)+becki (PBL)+*<br>*guntheri+microphyes+vandenburghi+vicina* | 0.113 (0.103–0.124) |
| *donfaustoi+porteri* | 0.287 (0.264–0.310) |
| *hoodensis+phantasticus+duncanensis+*<br>*chathamensis* | 0.144 (0.132–0.155) |

**Table S18.** Estimates of *gdi* for each taxon after the fifth round of sister taxa were collapsed. The *gdi* was calculated from the posterior distributions of *θ* and *τ* of the collapsed taxa in BPP, with prior *θ* ∼ IG(3, 0.001) and prior *τ* ∼ IG(3, 0.001).

| **Taxon** | **Median *gdi***<br>**(95% credible interval)** |
| --- | --- |
| *darwini+becki (PBR)+becki (PBL)+*<br>*guntheri+microphyes+vandenburghi+vicina+*<br>*donfaustoi+porteri* | 0.110 (0.102–0.118) |
| *hoodensis+ phantasticus+duncanensis+*<br>*chathamensis* | 0.142 (0.132–0.153) |

**Table S19.** Estimates of *gdi* for each taxon after the sixth round of sister taxa were collapsed. The *gdi* was calculated from the posterior distributions of *θ* and *τ* of the collapsed taxa in BPP, with prior *θ* ∼ IG(3, 0.001) and prior *τ* ∼ IG(3, 0.001).

| Taxon | Median *gdi*<br>(95% credible interval) |
| --- | --- |
| *darwini*+*becki* (*PBR*)+*becki* (*PBL*)+<br>*guntheri*+*microphyes*+*vandenburghi*+*vicina*+<br>*donfaustoi*+*porteri*+ *hoodensis*+<br>*phantasticus*+*duncanensis*+*chathamensis* | 0.988 (0.984–0.991) |
| *C. chilensis* | 0.92 (0.901–0.939) |

**Table S20:** Pairwise comparisons analyzed in PHRAPL across the two main clades of the phylogenetic tree, along with the wAIC for each of the three PHRAPL models. Models with a high Akaike weight (wAIC) > 0.9 for a model set are highlighted in gray. The cumulative sum of wAIC for all two species models is given for each comparison, with wAIC > 0.9 also highlighted in gray. Mean *gdi* < 0.2 are shown in bold.

| Taxon 1 | Taxon 2 | wAIC Model 1<br>2 species, no<br>gene flow | wAIC Model 2<br>2 species, constant<br>gene flow | wAIC Model 3<br>1 species | Sum wAIC of 2<br>Species<br>Models | Mean *gdi* |
| --- | --- | --- | --- | --- | --- | --- |
| **Within Saddleback Clade** |  |  |  |  |  |  |
| *duncanensis* | *hoodensis* | 0.984 | 0.016 | 0.000 | 1.000 | 0.387 |
| *chathamensis* | *hoodensis* | 0.465 | 0.535 | 0.000 | 1.000 | 0.384 |
| *chathamensis* | *duncanensis* | 0.722 | 0.278 | 0.000 | 1.000 | 0.217 |
| *hoodensis* | *phantasticus* | 0.539 | 0.461 | 0.000 | 1.000 | **0.179** |
| *duncanensis* | *phantasticus* | 0.022 | 0.978 | 0.000 | 1.000 | 0.319 |
| *chathamensis* | *phantasticus* | 0.000 | 1.000 | 0.000 | 1.000 | 0.319 |
| **Within Isabela Island (Domed Clade)** |  |  |  |  |  |  |
| *becki*(PBL) | *becki*(PBR) | 0.916 | 0.084 | 0.000 | 1.000 | **0.182** |
| *vicina* | *guntheri* | 0.227 | 0.773 | 0.000 | 1.000 | 0.447 |
| *vicina* | *vandenburghi* | 0.000 | 1.000 | 0.000 | 1.000 | 0.304 |
| *vicina* | *microphyes* | 0.005 | 0.995 | 0.000 | 1.000 | 0.345 |
| *vicina* | *becki*(PBL) | 0.127 | 0.873 | 0.000 | 1.000 | **0.179** |
| *vicina* | *becki*(PBR) | 0.000 | 1.000 | 0.000 | 1.000 | 0.307 |
| *guntheri* | *vandenburghi* | 0.001 | 0.999 | 0.000 | 1.000 | 0.455 |
| *guntheri* | *microphyes* | 0.000 | 1.000 | 0.000 | 1.000 | 0.460 |
| *guntheri* | *becki*(PBL) | 0.702 | 0.298 | 0.000 | 1.000 | 0.404 |
| *guntheri* | *becki*(PBR) | 0.000 | 1.000 | 0.000 | 1.000 | 0.536 |
| *vandenburghi* | *microphyes* | 0.790 | 0.210 | 0.000 | 1.000 | **0.180** |
| *vandenburghi* | *becki*(PBL) | 0.000 | 1.000 | 0.000 | 1.000 | 0.280 |
| *vandenburghi* | *becki*(PBR) | 0.000 | 1.000 | 0.000 | 1.000 | 0.316 |
| *microphyes* | *becki*(PBL) | 0.000 | 1.000 | 0.000 | 1.000 | 0.314 |
| *microphyes* | *becki*(PBR) | 0.006 | 0.994 | 0.000 | 1.000 | 0.344 |
| **Within Santa Cruz Island (Domed Clade)** |  |  |  |  |  |  |
| *donfaustoi* | *porteri* | 0.489 | 0.511 | 0.000 | 1.000 | **0.181** |
| **Santa Cruz Island with Isabela Island (Domed Clade)** |  |  |  |  |  |  |
| *donfaustoi* | *becki*(PBL) | 0.745 | 0.255 | 0.000 | 1.000 | 0.386 |
| *donfaustoi* | *becki*(PBR) | 0.004 | 0.996 | 0.000 | 1.000 | 0.458 |
| *donfaustoi* | *vandenburghi* | 0.665 | 0.335 | 0.000 | 1.000 | 0.386 |
| *donfaustoi* | *microphyes* | 0.000 | 1.000 | 0.000 | 1.000 | 0.484 |
| *donfaustoi* | *guntheri* | 0.000 | 1.000 | 0.000 | 1.000 | 0.276 |
| *donfaustoi* | *vicina* | 0.488 | 0.512 | 0.000 | 1.000 | 0.400 |
| *porteri* | *becki*(PBL) | 0.000 | 1.000 | 0.000 | 1.000 | 0.324 |
| *porteri* | *becki*(PBR) | 0.653 | 0.347 | 0.000 | 1.000 | 0.384 |
| *porteri* | *vandenburghi* | 0.665 | 0.335 | 0.000 | 1.000 | 0.386 |
| *porteri* | *microphyes* | 0.831 | 0.169 | 0.000 | 1.000 | 0.386 |
| *porteri* | *guntheri* | 0.000 | 1.000 | 0.000 | 1.000 | 0.281 |
| *porteri* | *vicina* | 0.000 | 1.000 | 0.000 | 1.000 | 0.344 |
| **Isabela and Santa Cruz Islands compared with Santiago Island (Domed Clade)** |  |  |  |  |  |  |
| *darwini* | *becki*(PBL) | 0.004 | 0.996 | 0.000 | 1.000 | 0.278 |
| *darwini* | *becki*(PBR) | 0.000 | 1.000 | 0.000 | 1.000 | 0.295 |
| *darwini* | *donfaustoi* | 0.597 | 0.403 | 0.000 | 1.000 | **0.181** |
| *darwini* | *porteri* | 0.203 | 0.797 | 0.000 | 1.00 | **0.181** |
| *darwini* | *guntheri* | 0.188 | 0.812 | 0.000 | 1.000 | **0.178** |
| *darwini* | *microphyes* | 0.000 | 1.000 | 0.000 | 1.000 | 0.375 |
| *darwini* | *vandenburghi* | 0.000 | 1.000 | 0.000 | 1.000 | 0.372 |
| *darwini* | *vicina* | 0.000 | 1.000 | 0.000 | 1.000 | 0.318 |

## References

Baele, G., Lemey, P., Bedford, T., Rambaut, A., Suchard, M. A., & Alekseyenko, A. V. (2012). Improving the accuracy of demographic and molecular clock model comparison while accommodating phylogenetic uncertainty. Molecular Biology and Evolution, 29(9), 2157–2167.

Benavides, E., Baum, R., Snell, H. M., Snell, H. L., & Sites, Jr, J. W. (2009). Island biogeography of Galápagos lava lizards (Tropiduridae: Microlophus): species diversity and colonization of the archipelago. Evolution, 63(6), 1606–1626.

Bouckaert, R., Heled, J., Kühnert, D., Vaughan, T., Wu, C. H., Xie, D., … & Drummond, A. J. (2014). BEAST 2: a software platform for Bayesian evolutionary analysis. PLoS Computational Biology, 10(4), e1003537.

Bryant, D., Bouckaert, R., Felsenstein, J., Rosenberg, N. A., & RoyChoudhury, A. (2012). Inferring species trees directly from biallelic genetic markers: bypassing gene trees in a full coalescent analysis. Molecular Biology and Evolution, 29(8), 1917–1932.

Caccone, A., Gibbs, J. P., Ketmaier, V., Suatoni, E., & Powell, J. R. (1999). Origin and evolutionary relationships of giant Galápagos tortoises. Proceedings of the National Academy of Sciences, 96(23), 13223–13228.

Caccone, A., Gentile, G., Gibbs, J. P., Fritts, T. H., Snell, H. L., Betts, J., & Powell, J. R. (2002). Phylogeography and history of giant Galápagos tortoises. Evolution, 56(10), 2052–2066.

Carstens, B. C., Pelletier, T. A., Reid, N. M., & Satler, J. D. (2013). How to fail at species delimitation. Molecular Ecology, 22(17), 4369–4383.

Chiari, Y. (2021). Morphology. In J. P. Gibbs, L. Cayot, and W. Tapia A (eds.). Galapagos Giant Tortoises. Academic Press.

de Queiroz, K. (1998). The general lineage concept of species, species criteria, and the process of speciation. In D. J. Howard and S. H. Berlocher (eds). Endless forms: species and speciation. Oxford University Press

de Queiroz, K. (2005a). Different species problems and their resolution. BioEssays, 27(12), 1263–1269.

de Queiroz, K. (2005b). A unified concept of species and its consequences for the future of taxonomy. Proceedings of the California Academy of Sciences.

de Queiroz, K. (2007). Species concepts and species delimitation. Systematic Biology, 56(6), 879–886.

Flouri, T., Jiao, X., Rannala, B., & Yang, Z. (2018). Species tree inference with BPP using genomic sequences and the multispecies coalescent. Molecular Biology and Evolution, 35(10), 2585–2593.

Garrick, R. C., Kajdacsi, B., Russello, M. A., Benavides, E., Hyseni, C., Gibbs, J. P., … & Caccone, A. (2015). Naturally rare versus newly rare: demographic inferences on two timescales inform conservation of Galápagos giant tortoises. Ecology and Evolution, 5(3), 676–694.

Gaughran, S. J., Quinzin, M. C., Miller, J. M., Garrick, R. C., Edwards, D. L., Russello, M. A., … & Caccone, A. (2018). Theory, practice, and conservation in the age of genomics: The Galápagos giant tortoise as a case study. Evolutionary Applications, 11(7), 1084–1093.

Geist, D. J., H. Snell, H. Snell, C. Goddard, and M. D. Kurz. (2014). A paleogeographic model of the Galápagos islands and biogeographical and evolutionary implications. Pp. 145-166 in K. S. Harpp, E. Mittelstaedt, N. d’Ozouville, and D. W. Graham (eds). The Galápagos: a natural laboratory for the earth science. John Wiley & Sons, Inc.

Giarla, T. C., Voss, R. S., & Jansa, S. A. (2014). Hidden diversity in the Andes: comparison of species delimitation methods in montane marsupials. Molecular Phylogenetics and Evolution, 70, 137–151.

Hey, J. (2006). On the failure of modern species concepts. Trends in Ecology & Evolution, 21(8), 447–450.

Hunter, E. A., Gibbs, J. P., Cayot, L. J., & Tapia, W. (2013). Equivalency of Galápagos giant tortoises used as ecological replacement species to restore ecosystem functions. Conservation Biology, 27(4), 701–709.

Jackson, N. D., Carstens, B. C., Morales, A. E., & O’Meara, B. C. (2017). Species delimitation with gene flow. Systematic Biology, 66(5), 799–812.

Jensen, E. L., Gaughran, S. J., Garrick, R. C., Russello, M. A., & Caccone, A. (2021). Demographic history and patterns of molecular evolution from whole genome sequencing in the radiation of Galapagos giant tortoises. Molecular Ecology, 30(23), 6325–6339.

Jensen, E. L., Gaughran, S. J., Fusco, N. A., Poulakakis, N., Tapia, W., Sevilla, C., … & Caccone, A. (2022). The Galapagos giant tortoise *Chelonoidis phantasticus* is not extinct. Communications Biology, 5(1), 546.

Kass, R. E., & Raftery, A. E. (1995). Bayes factors. Journal of the American Statistical Association, 90(430), 773–795.

Kehlmaier, C., Albury, N. A., Steadman, D. W., Graciá, E., Franz, R., & Fritz, U. (2021). Ancient mitogenomics elucidates diversity of extinct West Indian tortoises. Scientific Reports, 11(1), 3224.

Kimura, M., & Ohta, T. (1969). The average number of generations until fixation of a mutant gene in a finite population. Genetics, 61(3), 763.

Leaché, A. D., Zhu, T., Rannala, B., & Yang, Z. (2019). The spectre of too many species. Systematic Biology, 68(1), 168–181.

Leaché, A. D., Davis, H. R., Singhal, S., Fujita, M. K., Lahti, M. E., & Zamudio, K. R. (2021). Phylogenomic assessment of biodiversity using a reference-based taxonomy: an example with Horned Lizards (Phrynosoma). Frontiers in Ecology and Evolution, 9, 678110.

Li, H., Handsaker, B., Wysoker, A., Fennell, T., Ruan, J., Homer, N., … & 1000 Genome Project Data Processing Subgroup. (2009). The sequence alignment/map format and SAMtools. Bioinformatics, 25(16), 2078–2079.

Loire, E., Chiari, Y., Bernard, A., Cahais, V., Romiguier, J., Nabholz, B., … & Galtier, N. (2013). Population genomics of the endangered giant Galápagos tortoise. Genome Biology, 14, 1–11.

Mallet, J. (1995). A species definition for the modern synthesis. Trends in Ecology & Evolution, 10(7), 294–299.

Mays Jr, H. L., Oehler, D. A., Morrison, K. W., Morales, A. E., Lycans, A., Perdue, J., … & Weakley, L. A. (2019). Phylogeography, population structure, and species delimitation in rockhopper penguins (*Eudyptes chrysocome* and *Eudyptes moseleyi*). Journal of Heredity, 110(7), 801–817.

Mayden, R. L. (1997). A hierarchy of species concepts: the denouement in the saga of the species problem. pp. 381–423 in M. F. Claridge, H. A. Dawah & M. R. Wilson (eds.). Species: the Units of Diversity. Chapman & Hall.

Parent, C. E., Caccone, A., & Petren, K. (2008). Colonization and diversification of Galápagos terrestrial fauna: a phylogenetic and biogeographical synthesis. Philosophical Transactions of the Royal Society B: Biological Sciences, 363(1508), 3347–3361.

Poulakakis, N., Glaberman, S., Russello, M., Beheregaray, L. B., Ciofi, C., Powell, J. R., & Caccone, A. (2008). Historical DNA analysis reveals living descendants of an extinct species of Galápagos tortoise. Proceedings of the National Academy of Sciences, 105(40), 15464–15469.

Poulakakis, N., Russello, M., Geist, D., & Caccone, A. (2012). Unravelling the peculiarities of island life: vicariance, dispersal and the diversification of the extinct and extant giant Galápagos tortoises. Molecular Ecology, 21(1), 160–173.

Poulakakis, N., Edwards, D. L., Chiari, Y., Garrick, R. C., Russello, M. A., Benavides, E., … & Caccone, A. (2015). Description of a new Galápagos giant tortoise species (Chelonoidis; Testudines: Testudinidae) from Cerro Fatal on Santa Cruz Island. PLoS One, 10(10), e0138779.

Poulakakis, N., Miller, J. M., Jensen, E. L., Beheregaray, L. B., Russello, M. A., Glaberman, S., … & Caccone, A. (2020). Colonization history of Galapagos giant tortoises: Insights from mitogenomes support the progression rule. Journal of Zoological Systematics and Evolutionary Research, 58(4), 1262–1275.

Pritchard, P. C. (1996). The Galápagos tortoises: nomenclature and survival status. Chelonian Research Foundation.

Quesada, V., Freitas-Rodríguez, S., Miller, J., Pérez-Silva, J. G., Jiang, Z. F., Tapia, W., … & López-Otín, C. (2019). Giant tortoise genomes provide insights into longevity and age-related disease. Nature ecology & evolution, 3(1), 87–95.

R Core Team (2022). R: A language and environment for statistical computing. R Foundation for Statistical Computing, Vienna, Austria. URL https://www.R-project.org/.

Reilly, S. B., Stubbs, A. L., Arida, E., Karin, B. R., Arifin, U., Kaiser, H., … & McGuire, J. A. (2022). Phylogenomic analysis reveals dispersal-driven speciation and divergence with gene flow in Lesser Sunda flying lizards (Genus *Draco*). Systematic Biology, 71(1), 221–241.

Rhodin, A.G.J., Iverson, J.B., Bour, R., Fritz, U., Georges, A., Shaffer, H.B., and van Dijk, P.P. (2021). Turtles of the World: Annotated Checklist and Atlas of Taxonomy, Synonymy, Distribution, and Conservation Status (9th Ed.). In Rhodin, A.G.J., Iverson, J.B., van Dijk, P.P., Stanford, C.B., Goode, E.V., Buhlmann, K.A., and Mittermeier, R.A. (Eds.). Conservation Biology of Freshwater Turtles and Tortoises: A Compilation Project of the IUCN/SSC Tortoise and Freshwater Turtle Specialist Group. Chelonian Research Monographs 8:1–472. doi:10.3854/crm.8.checklist.atlas.v9.2021.

Ruffley, M., Smith, M. L., Espíndola, A., Carstens, B. C., Sullivan, J., & Tank, D. C. (2018). Combining allele frequency and tree-based approaches improves phylogeographic inference from natural history collections. Molecular Ecology, 27(4), 1012–1024.

Russello, M. A., Glaberman, S., Gibbs, J. P., Marquez, C., Powell, J. R., & Caccone, A. (2005). A cryptic taxon of Galápagos tortoise in conservation peril. Biology Letters, 1(3), 287–290.

Schafer, S. F., & Krekorian, C. N. (1983). Agonistic behavior of the Galapagos tortoise, *Geochelone elephantopus*, with emphasis on its relationship to saddle-backed shell shape. Herpetologica, 448–456.

Smith, M. L., & Carstens, B. C. (2020). Process-based species delimitation leads to identification of more biologically relevant species. Evolution, 74(2), 216–229.

Stamatakis A. (2014). RAxML version 8: a tool for phylogenetic analysis and post-analysis of large phylogenies. Bioinformatics. 30(9):1312–3

Yang, Z., & Rannala, B. (2010). Bayesian species delimitation using multilocus sequence data. Proceedings of the National Academy of Sciences, 107(20), 9264–9269.

## Supplemental References

Austin, C. C., Rittmeyer, E. N., Richards, S. J., & Zug, G. R. (2010). Phylogeny, historical biogeography and body size evolution in Pacific Island Crocodile skinks Tribolonotus (Squamata; Scincidae). Molecular Phylogenetics and Evolution, 57(1), 227–236.

Bergeron, L. A., Besenbacher, S., Zheng, J., Li, P., Bertelsen, M. F., Quintard, B., … & Zhang, G. (2023). Evolution of the germline mutation rate across vertebrates. Nature, 615(7951), 285–291.

Ciofi, C., Milinkovitch, M. C., Gibbs, J. P., Caccone, A., & Powell, J. R. (2002). Microsatellite analysis of genetic divergence among populations of giant Galápagos tortoises. Molecular Ecology, 11(11), 2265–2283.

Ciofi, C., Wilson, G. A., Beheregaray, L. B., Marquez, C., Gibbs, J. P., Tapia, W., … & Powell, J. R. (2006). Phylogeographic history and gene flow among giant Galápagos tortoises on southern Isabela Island. Genetics, 172(3), 1727–1744.

Drummond, A. J., & Rambaut, A. (2007). BEAST: Bayesian evolutionary analysis by sampling trees. BMC Evolutionary Biology, 7, 214.

Garrick, R. C., Benavides, E., Russello, M. A., Hyseni, C., Edwards, D. L., Gibbs, J. P., … & Caccone, A. (2014). Lineage fusion in Galápagos giant tortoises. Molecular Ecology, 23(21), 5276–5290.

Hedrick, P. W. (2009). Genetics of populations. Jones & Bartlett Publishers.

Iannucci, A., Benazzo, A., Natali, C., Arida, E. A., Zein, M. S. A., Jessop, T. S., … & Ciofi, C. (2021). Population structure, genomic diversity and demographic history of Komodo dragons inferred from whole-genome sequencing. Molecular Ecology, 30(23), 6309–6324.

Li, H. (2013). Aligning sequence reads, clone sequences and assembly contigs with BWA-MEM. arXiv preprint arXiv:1303.3997.

Li, H., Handsaker, B., Wysoker, A., Fennell, T., Ruan, J., Homer, N., … & 1000 Genome Project Data Processing Subgroup. (2009). The sequence alignment/map format and SAMtools. Bioinformatics, 25(16), 2078-2079.

Miller, J. M., Quinzin, M. C., Edwards, D. L., Eaton, D. A., Jensen, E. L., Russello, M. A., … & Caccone, A. (2018). Genome-wide assessment of diversity and divergence among extant Galapagos giant tortoise species. Journal of Heredity, 109(6), 611–619.

Quesada, V., Freitas-Rodríguez, S., Miller, J., Pérez-Silva, J. G., Jiang, Z. F., Tapia, W., … & López-Otín, C. (2019). Giant tortoise genomes provide insights into longevity and age-related disease. Nature Ecology & Evolution, 3(1), 87–95.

Rambaut, A., Drummond, A. J., Xie, D., Baele, G., & Suchard, M. A. (2018). Posterior summarization in Bayesian phylogenetics using Tracer 1.7. Systematic Biology, 67(5), 901–904.

Rannala, B. and Yang, Z. (2020). Species Delimitation. In Scornavacca, C., Delsuc, F., and Galtier, N. (eds.). pp. 5.5:1–5.5:18 in Phylogenetics in the Genomic Era. No commercial publisher | Authors open access book. https://hal.inria.fr/PGE

Stephens, M., Smith, N. J., & Donnelly, P. (2001). A new statistical method for haplotype reconstruction from population data. The American Journal of Human Genetics, 68(4), 978–989.

Stephens, M., & Scheet, P. (2005). Accounting for decay of linkage disequilibrium in haplotype inference and missing-data imputation. The American Journal of Human Genetics, 76(3), 449–462.

